# Combining antibiotics with antivirulence compounds can have synergistic effects and reverse selection for antibiotic resistance in *Pseudomonas aeruginosa*

**DOI:** 10.1101/861799

**Authors:** Chiara Rezzoagli, Martina Archetti, Ingrid Mignot, Michael Baumgartner, Rolf Kümmerli

## Abstract

Antibiotics are losing efficacy due to the rapid evolution and spread of resistance. Treatments targeting bacterial virulence factors have been considered as alternatives because they target virulence instead of pathogen viability, and should therefore exert weaker selection for resistance than conventional antibiotics. However, antivirulence treatments rarely clear infections, which compromises their clinical applications. Here, we explore the potential of combining antivirulence drugs with antibiotics against the opportunistic human pathogen *Pseudomonas aeruginosa*. We combined two antivirulence compounds (gallium, a siderophore-quencher, and furanone C-30, a quorum sensing-inhibitor) together with four clinically relevant antibiotics (ciprofloxacin, colistin, meropenem, tobramycin) in 9×9 drug concentration matrices. We found that drug-interaction patterns were concentration dependent, with promising levels of synergies occurring at intermediate drug concentrations for certain drug pairs. We then tested whether antivirulence compounds are potent adjuvants, especially when treating antibiotic resistant clones. We found that the addition of antivirulence compounds to antibiotics could restore growth inhibition for most antibiotic resistant clones, and even abrogate or reverse selection for resistance in five drug combination cases. Molecular analyses suggest that selection against resistant clones occurs when resistance mechanisms involve restoration of protein synthesis, but not when efflux pumps are upregulated. Altogether, our work provides a first systematic analysis of antivirulence-antibiotic combinatorial treatments and suggests that such combinations have a high potential to be both effective in treating infections and in limiting the spread of antibiotic resistance.

## Introduction

Scientists together with the World Health Organization (WHO) forecast that the rapid evolution and spread of antibiotic resistant bacteria will lead to a world-wide medical crisis [1–3]. Already today, the effective treatment of an increasing number of infectious diseases has become difficult in many cases [4,5]. To avert the crisis, novel innovative approaches that are both effective against pathogens and robust to the emergence and spread of resistance are urgently needed [6,7]. One such approach involves the use of compounds that disarm rather than kill bacteria. These so-called ‘antivirulence’ treatments should exert weaker selection for resistance compared to classical antibiotics because they simply disable virulence factors but are not supposed to affect pathogen viability [8–10]. However, a downside of antivirulence approaches is that the infection will not necessarily be cleared. This could be particularly problematic for immuno-compromised patients (AIDS, cancer, cystic fibrosis and intensive-care unit patients), whose immune system is conceivably too weak to clear even disarmed pathogens.

One way to circumvent this problem is to combine antivirulence compounds with antibiotics to benefit from both virulence suppression and effective pathogen removal [6,11]. While a few studies have already considered such combinatorial treatments [12–18], we currently have no comprehensive understanding of how different types of antibiotics and antivirulence drugs interact, whether interactions are predominantly synergistic or antagonistic, and how combinatorial treatments affect growth and the spread of antibiotic resistant strains. Such knowledge is, however, essential if such therapies are supposed to make their way into the clinics, as drug interactions and their effects on antibiotic resistance evolution will determine both the efficacy and sustainability of treatments. Here, we tackle these open issues by combining four different classes of antibiotics with two antivirulence compounds as treatments against the opportunistic human pathogen *Pseudomonas aeruginosa*, to test the nature of drug interactions, and the usefulness of antivirulence compounds as adjuvants to combat antibiotic sensitive and resistant strains.

*P. aeruginosa* is one of the ESKAPE pathogens with multi-drug resistant strains spreading worldwide and infections becoming increasingly difficult to treat [19,20]. In addition to its clinical relevance, *P. aeruginosa* has become a model system for antivirulence research [21]. Several antivirulence compounds targeting either the species’ quorum sensing (QS) [22–24] or siderophore-mediated iron uptake systems [25–28] have been proposed. While QS is a cell-to-cell communication system that controls the expression of multiple virulence factors including exo-proteases, biosurfactants and toxins, siderophores are secondary metabolites important for the scavenging of iron from host tissue. For our experiments, we chose antivirulence compounds that target these two different virulence mechanisms: furanone C-30, an inhibitor of the LasR QS-system [12,22], and gallium, targeting the iron-scavenging pyoverdine and iron-metabolism [25,27,29–33].

Furanone C-30 is a synthetic, brominated furanone, which includes the same lactone ring present in the acylhomoserine lactones (AHL) QS molecules of *P. aeruginosa* [34]. As a consequence, it can disrupt QS-based communication by antagonistically competing with the AHLs molecules for binding to the main QS (LasR) receptor in a concentration dependent manner [12,35]. Gallium is an iron mimic whose ionic radius and coordination chemistry is comparable to ferric iron, although its redox properties are different. Specifically, gallium cannot be reduced and thereby irreversibly binds to siderophores and hinders siderophore-mediated iron uptake [25,27]. Pyoverdine is controlled in complex ways involving information on cell density and feedbacks from iron uptake rates [36,37]. Gallium interferes with these regulatory circuits, whereby cells intermittently upregulate pyoverdine production at low gallium concentrations to compensate for the increased level of iron limitation, and down-regulate pyoverdine production at higher gallium concentrations because iron uptake via this pathway is unsuccessful [27]. Importantly, gallium drastically reduces pyoverdine availability at the population level in a concentration-dependent manner [25,38].

We combined the two antivirulence compounds with four clinically relevant antibiotics (ciprofloxacin, colistin, meropenem, and tobramycin), which are widely used against *P. aeruginosa* [39]. In a first step, we measured treatment effects on bacterial growth and virulence factor production for all eight drug combinations for 81 concentration combinations each. In a second step, we applied the Bliss independence model to calculate the degree of synergy or antagonism to obtain comprehensive interaction maps for all combinations both for growth and virulence factor production. Next, we selected for antibiotic resistant clones and tested whether the addition of antivirulence compounds as adjuvants restores growth inhibition and affects selection for antibiotic resistance. Finally, we sequenced the genomes of the evolved antibiotic resistant clones to understand the genetic basis that drive the observed effects on pathogen growth and selection for or against antibiotic resistance.

## Results

### PAO1 dose-response curves to antibiotics and antivirulence compounds

In a first experiment, we determined the dose-response curve of PAO1 to each of the four antibiotics (Fig 1) and the two anti-virulence compounds (Fig 2) in our experimental media. We found that the dose-response curves for antibiotics followed sigmoid functions (Fig 1), characterised by (i) a low antibiotic concentration range that did not inhibit bacterial growth, (ii) an intermediate antibiotic concentration range that significantly reduced bacterial growth, and (iii) a high antibiotic concentration range that completely stalled bacterial growth. Since antibiotics curbed growth, they congruently also reduced the availability of virulence factors at the population level (S1 Fig).

**Fig 1.**
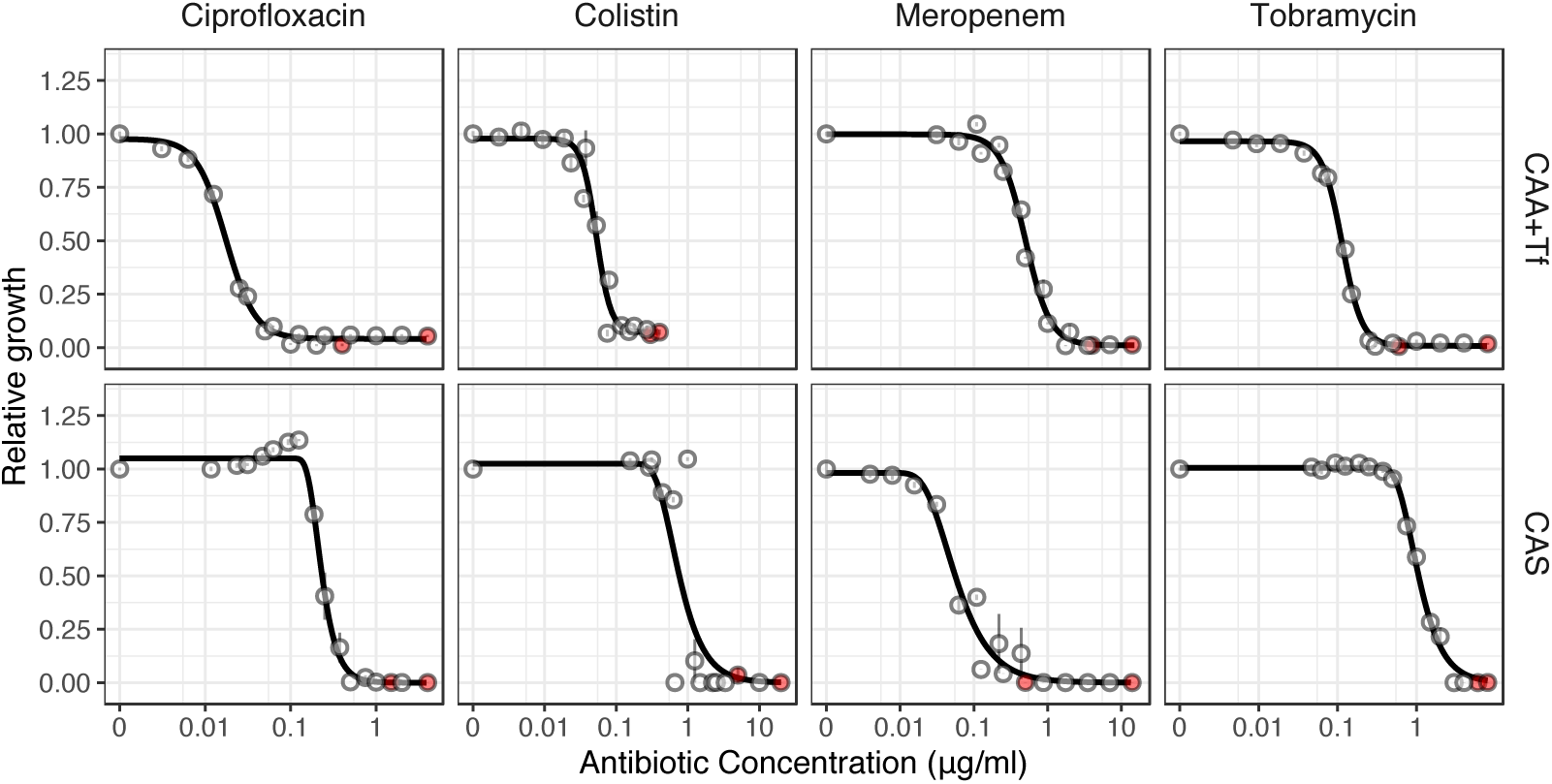
Antibiotic dose response curves for *P. aeruginosa* PAO1. We exposed PAO1 to all four antibiotics in two experimental media: CAA+Tf (iron-limited casamino acids medium with transferrin) and CAS (casein medium). Except for meropenem, higher concentrations of antibiotics were required to inhibit PAO1 in CAS compared to CAA+Tf. Dots show means ± standard error across six replicates. All data are scaled relative to the drug-free treatment. Data stem from two independent experiments using different dilution series. The red dots indicate the highest concentration used for the respective experiments, from which 7-serial dilution steps were tested. Curves were fitted with either log-logistic functions (in CAA+Tf) or with three-parameter Weibull functions (in CAS).

**Fig 2.**
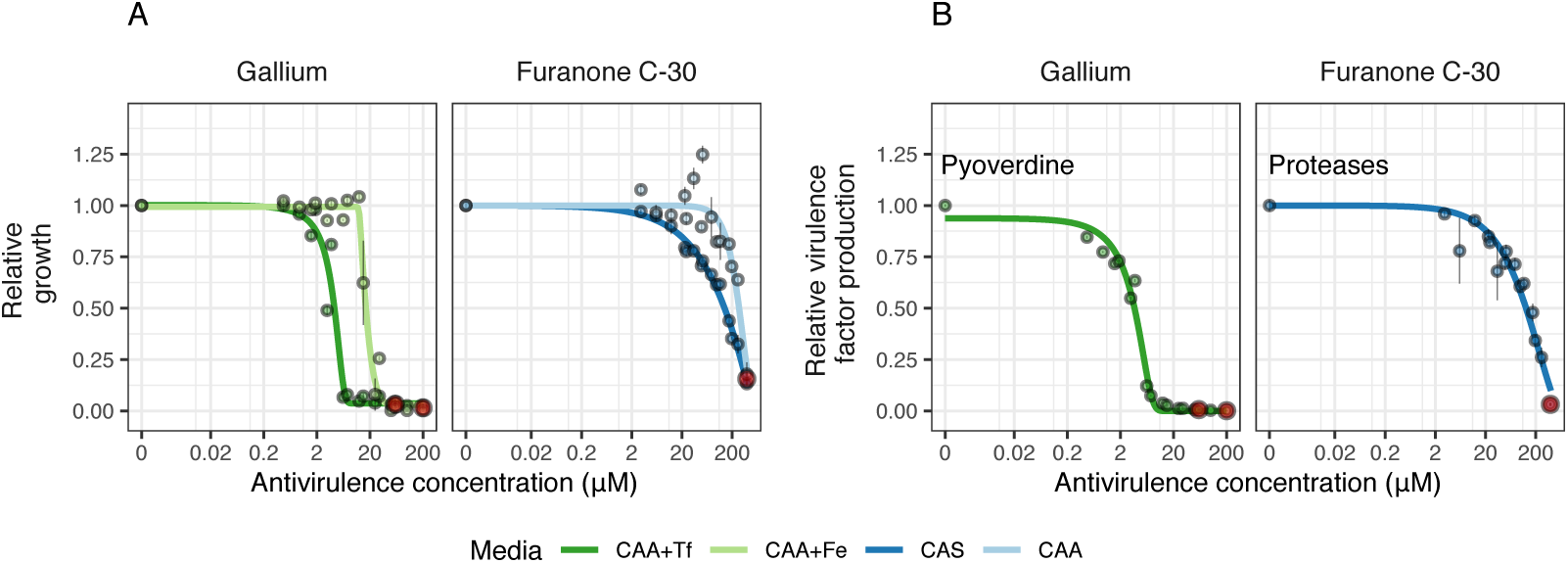
Antivirulence dose response curves for *P. aeruginosa* PAO1 (growth and virulence factor production). We exposed PAO1 to the antivirulence compounds gallium (inhibiting pyoverdine-mediated iron uptake) and furanone C-30 (blocking quorum sensing response including protease production) both in media where the targeted virulence factors are expressed and required (iron-limited CAA+Tf medium for gallium and CAS medium for furanone) and in control media where the targeted virulence factors are not required (iron-supplemented CAA+Fe medium for gallium and protein digested CAA for furanone). (**A**) Dose-response curves for growth show that both antivirulence compounds reduced bacterial growth, but more so in media where the targeted virulence factor is expressed. This demonstrates that there is a concentration window where the antivirulence compounds have no toxic effects on bacterial cells and just limit growth due to virulence factor quenching. (**B**) Dose-response curves for virulence factor production show that gallium and furanone C-30 effectively inhibit pyoverdine and protease production, respectively, in a concentration-dependent manner. Dots show means ± standard errors across six replicates. All data are scaled relative to the drug-free treatment. Data stem from two independent experiments using different dilution series. The red dots indicate the highest concentration used for the respective experiments, from which 7-serial dilution steps were tested. Curves were fitted with either log-logistic functions (in CAA+Tf) or with three-parameter Weibull functions (in CAS).

Similarly shaped dose-response curves for growth (Fig 2A) and virulence factor production (Fig 2B) were obtained for gallium (quenching pyoverdine) and furanone C-30 (inhibiting protease production) in the respective media where the two virulence factors are important for growth. Under such conditions, the reduced availability of virulence factors immediately feeds back on growth by inducing iron starvation (gallium) or the inability to degrade proteins (furanone). Crucially, the dose-response curves for growth shifted to the right (extending phase (i)) when we repeated the experiment in media, where the virulence factors are not needed for growth (i.e. iron-rich media for pyoverdine, and protein digest media for proteases). This shows that there is a window of concentrations where growth inhibition is caused by virulence factor quenching alone. Conversely, high concentrations of antivirulence compounds seem to have additional off-target effects reducing growth.

### Interaction maps of antibiotic-antivirulence drug combinations

#### General patterns

From the dose-response curves, we chose 9 concentrations for each drug to cover the entire trajectory, from no to intermediate to high growth inhibition. We then combined antibiotics with antivirulence compounds in a 9×9 concentration matrix and measured the dose-response curve for every single drug combination for both growth and virulence factor production (Fig 3). At the qualitative level, independent drug effects would cause a symmetrical downshift of the dose-response curve with higher antivirulence compound concentrations supplemented. We indeed noticed symmetrical downshifts for many dose-response curves (Fig 3), but there were also clear cases of non-symmetrical shifts, indicating synergy or antagonism between drugs. We then used the Bliss model, representing the adequate model for drugs with independent modes of actions, to quantify these effects. We found patterns of synergy and antagonism for both growth and virulence factor inhibition across the concentration matrices for all drug combinations (Fig 4), with many of these interactions being significant (S2 Fig).

**Fig 3.**
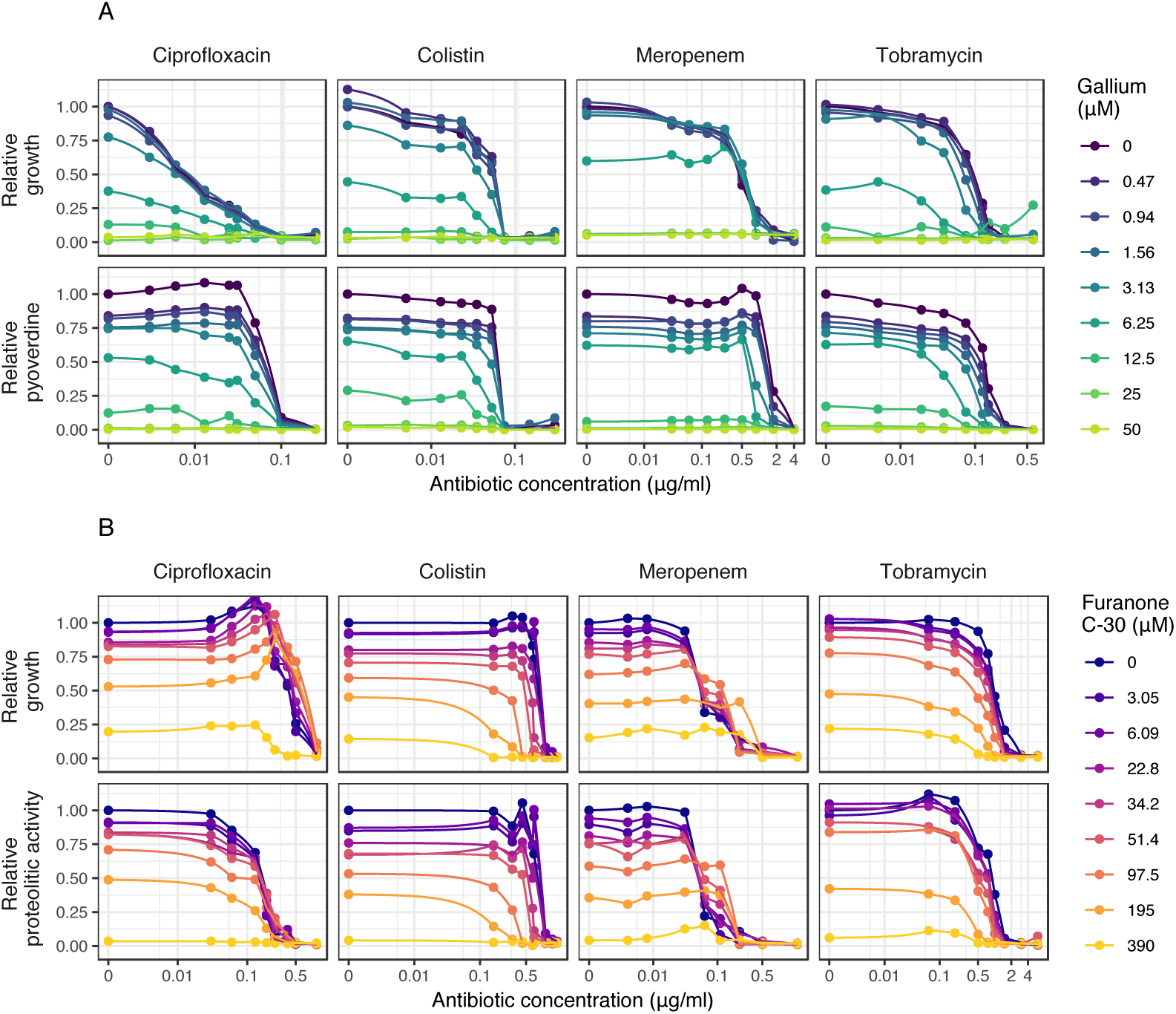
Dose response curves for *P. aeruginosa* PAO1 under antibiotic-antivirulence combination treatments. Dose-response curves for growth and virulence factor production for PAO1 were assessed for 9×9 drug concentration matrixes involving the four antibiotics combined with either gallium (**A**) or furanone C-30 (**B**). Experiments were carried out in media where the corresponding virulence factors are required for growth (pyoverdine: CAA+Tf; protease: CAS). Growth and virulence factor production were measured after 48 hours. All values are scaled relative to the untreated control, and data points show the mean across 12 replicates from two independent experiments. We used spline functions to fit the dose-response curves.

**Fig 4:**
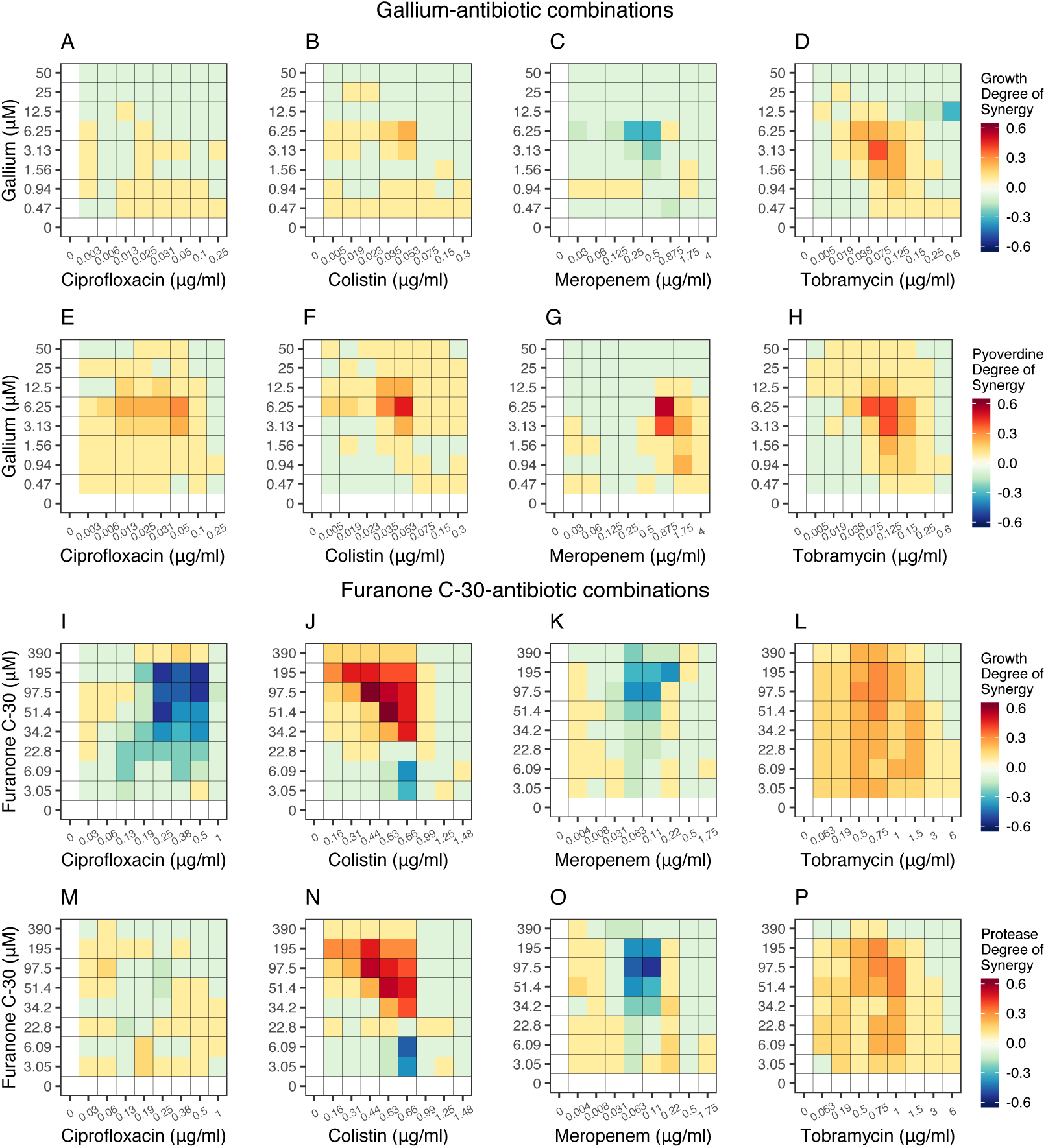
Drug interaction heatmaps for antibiotic-antivirulence combination treatments. We used the Bliss independence model to calculate the degree of synergy for every single drug combination with regard to growth suppression and virulence factor quenching shown in Fig 3. Heatmaps, depicting variation in drug interactions ranging from antagonism (blue) to synergy (red), are shown for gallium-antibiotic combinations (**A-D** for growth; **E-H** for pyoverdine production) and furanone-antibiotic combinations (**I-L** for growth; **M-P** for protease production). All calculations are based on 12 replicates from two independent experiments.

#### Gallium-antibiotic combinations

Gallium combined with ciprofloxacin or colistin had mostly independent effects on bacterial growth (i.e. weak or no synergy/antagonism) (Fig 4A+B). With regard to the inhibition of pyoverdine production, both drug combinations showed significant levels of synergy at intermediate drug concentrations (Fig 4E+F, S2 Fig). For gallium-meropenem combinations, we observed mostly independent interactions for growth and pyoverdine inhibition, with small hotspots of antagonism (for growth) and synergy (for siderophore inhibition) existing at intermediate drug concentrations (Fig 4C+G). Finally, for gallium-tobramycin combinations there were relatively strong significant synergistic interactions for both growth and pyoverdine inhibition at intermediate drug concentrations (Fig 4D+H, S2 Fig). Interestingly, we observed synergy with regard to pyoverdine inhibition for all drug combinations, indicating that the combination of low cell density induced by the antibiotics and gallium-mediated pyoverdine quenching is a successful strategy to repress this virulence factor.

#### Furanone-antibiotic combinations

For furanone-ciprofloxacin combinations, we found relatively strong significant antagonistic interactions with regard to growth inhibition (Fig 4I), whereas effects on protease inhibition were mostly independent (Fig 4M). In contrast, for furanone-colistin combinations we observed strong and significant synergistic drug interactions especially for intermediate and higher concentrations of the antivirulence compound for growth and protease inhibition (Fig 4J+N, S2 Fig). Furanone-meropenem, on the other hand, interacted mostly antagonistically with regard to growth and protease inhibition (Fig 4K+O). Conversely, for furanone-tobramycin combinations there were pervasive significant patterns of synergy across the entire drug combination range for growth and virulence factor inhibition (Fig 4L+P, S2 Fig).

#### Do the degrees of synergy for growth and virulence factor inhibition correlate?

As the combination treatments affect both growth and virulence factor production, we examined whether the degrees of synergy correlate between the two traits (S3 Fig). For gallium-antibiotic combinations, we found no correlations for ciprofloxacin and meropenem, but positive associations for colistin and tobramycin (Pearson correlation coefficient; ciprofloxacin: r = 0.09, t_79_ = 0.85, p = 0.394; colistin: r = 0.69, t_79_ = 8.51, p < 0.001; meropenem: r = 0.17, t_79_ = 1.53, p = 0.130; tobramycin: r = 0.58, t_79_ = 6.39, p < 0.001). For furanone-antibiotic combinations, there were strong positive correlations between the levels of synergy for the two traits for all drug combinations (ciprofloxacin: r = 0.34, t_79_ = 3.22, p = 0.002; colistin: r = 0.96, t_79_ = 32.50, p < 0.001; meropenem: r = 0.87, t_79_ = 15.48, p < 0.001; tobramycin: r = 0.75, t_79_ = 10.16, p < 0.001).

### Antibiotic resistance can lead to collateral sensitivity and cross-resistance to antivirulence compounds

In a next step, we asked whether antivirulence compounds could be used as adjuvants to suppress the growth of antibiotic resistant clones. To address this question, we first experimentally selected, isolated, and analyzed antibiotic resistant clones (AtbR). We aimed for one clone per antibiotic and medium. In the end, we examined seven clones, as only one selection line survived the ciprofloxacin selection regime (see methods for details). We then assessed the dose-response curve of these clones for the respective antibiotics to confirm and quantify their level of resistance (S4 Fig). We further established the dose-response curves of all AtbR clones for the two antivirulence compounds to test for collateral sensitivity and cross-resistance. Here, we compared the IC50 (half maximal inhibitory concentration) values between the AtbR and wildtype strain (Fig S5), and found evidence for weak but significant collateral sensitivity between colistin and gallium and relatively strong collateral sensitivity between tobramycin and furanone. Conversely, we found that resistance to ciprofloxacin, colistin and also to some extent to meropenem can confer cross-resistance to furanone (all statistical analyses are shown in S5 Fig).

Based on these experiments, we picked two concentrations for gallium (low: 1.56 µM; intermediate: 6.25 µM) and furanone (low: 6.3 µM; intermediate: 22.8 µM) as adjuvants in combination with antibiotics to test whether antivirulence compounds can restore growth inhibition of and alter selection for AtbR clones.

### Antivirulence compounds as adjuvants can restore growth inhibition of antibiotic resistant strains

In a first set of experiments, we subjected all AtbR clones and the antibiotic-sensitive wildtype to antivirulence treatments alone and to combination treatments with antibiotics (Fig 5). For gallium, we observed that the wildtype and AtbR clones responded almost identically to the antivirulence compounds in the absence of antibiotics, showing that AtbR clones are still sensitive to gallium (Fig 5A). When treated with antibiotics, we observed that the addition of anti-virulence compounds consistently reduced growth of all AtbR clones (Fig 5A). The level of synergy between drugs is similar for wildtype and AtbR clones, with the effects being close to zero (varying between weak antagonism and weak synergy) in most cases. Altogether, these findings indicate that gallium acts independently to all the tested antibiotics and is still able induce iron starvation and thus reduce growth in AtbR clones. Important to note is that gallium has a stronger growth inhibitory effect in this experiment compared to the dose-response curve analysis (S5 Fig, collected after 48 hours), because this experiment run for 24 hours only.

**Fig 5:**
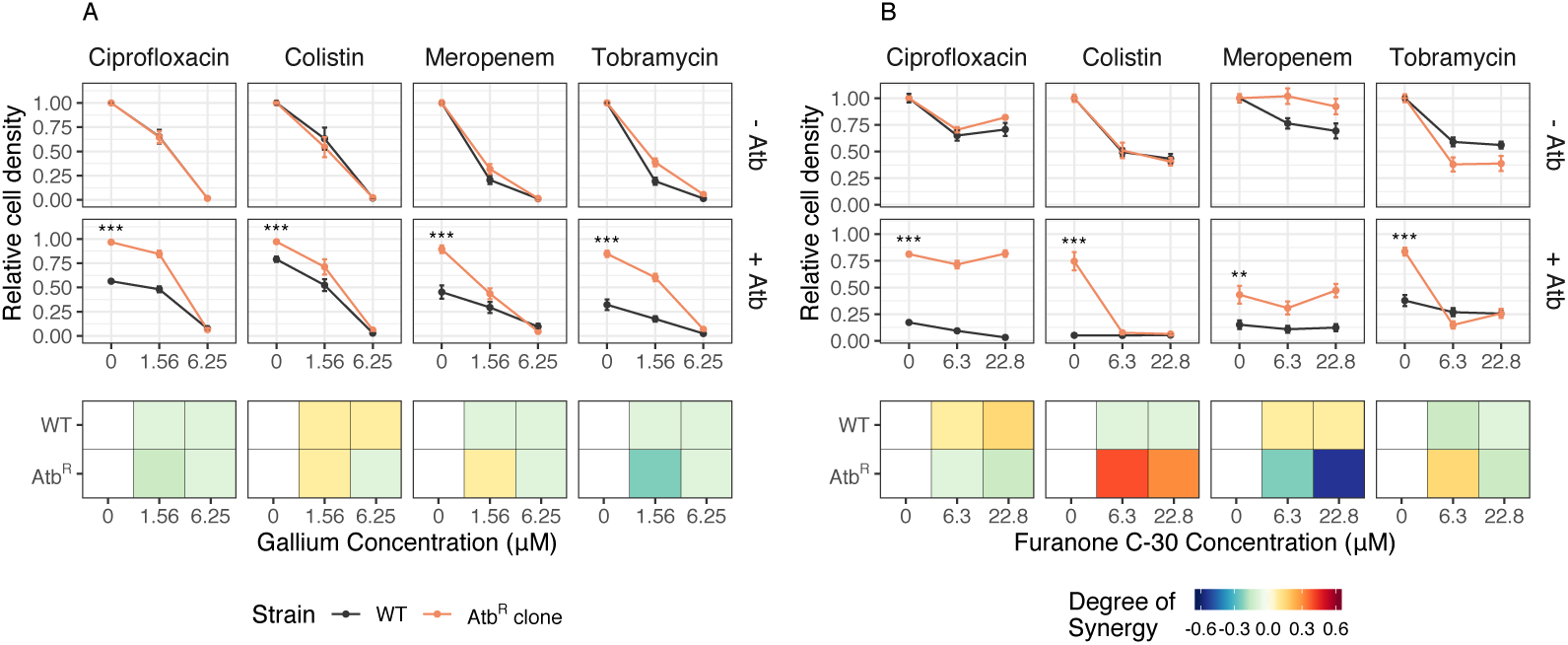
Effect of combination treatment on growth of antibiotic resistant clones. Test of whether the addition of gallium (**A**) or furanone (**B**) can restore growth suppression in antibiotic resistant clones (AtbR, in orange) relative to the susceptible wildtype (WT, in black). Under antibiotic treatment and in the absence of antivirulence compounds, all AtbR strains grew significantly better than the WT (two-sample t-tests, - 26.63 ≤ *t*_21-40_ ≤ −3.03, p < 0.01 for all treatment combinations; n.s. = non-significant; * p <0.05; ** p <0.01; *** p <0.001), a result that holds for both scaled (as shown above) and absolute growth. In the presence of antivirulence compounds (upper series of panels without antibiotics (-Atb); lower series of panels with antibiotics (+Atb)), growth suppression was restored in six out of eight cases. The exceptions were the ciprofloxacin-furanone and meropenem-furanone combinations. The bottom series of panels shows the degree of drug synergy for the wildtype and the AtbR clones. All cell density values (measured with flow cytometry as number of events detected in 5 μl of culture, after 24 hours) are scaled relative to the untreated control. All data are shown as means ± standard errors across a minimum of 16 replicates from 4 to 6 independent experiments.

For furanone, the patterns were more diverse (Fig 5B). There were two cases of full cross-resistance (ciprofloxacin and meropenem), where the addition of furanone no longer had any effect on bacterial growth. In these cases, we observed a change from weak drug synergy (for the wildtype) to strong drug antagonism (for the AtbR clones). In contrast, we found a strong shift from weak antagonism (for the wildtype) to strong drug synergy (for the AtbR clone) when furanone was combined with colistin. In this case, furanone re-potentiated the antibiotic. Note that we initially also observed a certain level of cross-resistance for this drug combination (S5 Fig), however only at much higher furanone concentrations than those used here. Finally, the pattern between tobramycin and furanone was driven by collateral sensitivity, where the addition of furanone to the AtbR clone completely restored growth inhibition.

### Antivirulence as adjuvants can abrogate or reverse selection for antibiotic resistance

We then investigated whether antivirulence compounds alone or in combination with antibiotics can influence the spread of AtbR clones in mixed populations with susceptible wildtype cells (Fig 6). First, we competed the AtbR clones against the susceptible wildtype in the absence of any treatment and observed that AtbR clones consistently lost the competitions (one sample t-tests, −13.50 ≤ *t*_15-26_ ≤ −2.62, p ≤ 0.050 for all comparisons). This confirms that antibiotic resistance is costly and is selected against in the absence of treatment. We then added the antivirulence drug alone using the same concentrations as for the monoculture experiments (Fig 5). We found that the AtbR clones did not experience a selective advantage in 14 out 16 cases, with the exception being one colistin resistant clone that slightly increased in frequency (Fig 6). This analysis shows that the cost of AtbR resistance is largely maintained in the presence of antivirulence compounds. Next, we exposed the mixed cultures to the antibiotics alone and observed that, as expected, AtbR clones always experienced a significant fitness advantage under treatment (one sample t-test, 4.54 ≤ *t*_19-26_ ≤ 13.41, p < 0.001 for all combinations).

**Fig 6:**
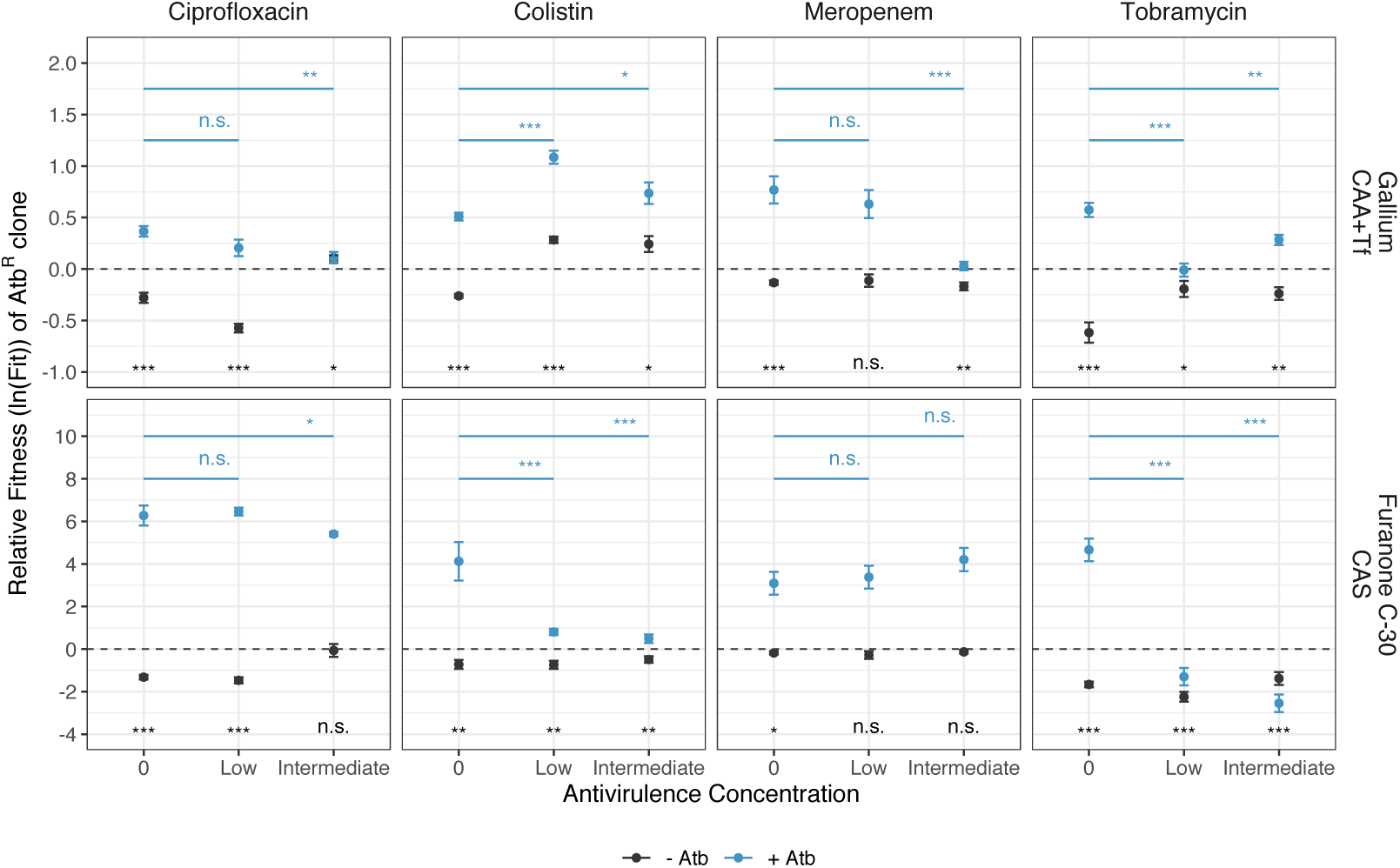
Effect of combination treatment on the relative fitness of antibiotic resistant clones. Test of whether antivirulence compounds alone or in combination with antibiotics can abrogate or revert selection for antibiotic resistance. All AtbR clones were competed against the susceptible wildtype (WT) for 24 hours, starting at a 1:9 ratio. The dashed lines denote fitness parity, where none of the competing strains has a fitness advantage. In the absence of any treatment, all AtbR clones showed a fitness disadvantage (fitness values < 0) compared to the WT, demonstrating the cost of resistance. When treated with antivirulence compounds alone, the AtbR clones did not experience any selective advantage in 14 out of 16 cases (exception: colistin-gallium combinations). When treated with antibiotics alone, all AtbR clones experienced significant fitness advantages (fitness values > 0), as expected. When antivirulence compounds were added as adjuvants to antibiotics, the fitness advantage of AtbR clones was reduced, abrogated or reversed for six out of eight drug combinations. All data are shown as means ± standard errors across a minimum of 16 replicates from 4 to 7 independent experiments. Significance levels are based on t-tests or ANOVAs: n.s. = non-significant; * p <0.05; ** p <0.01; *** p <0.001. See Table S1 for full details on the statistical analyses.

When combining antivirulence compounds with antibiotics, three different relative fitness patterns emerged for AtbR clones. In three cases (colistin-gallium, ciprofloxacin-furanone and meropenem-furanone), AtbR clones experienced large fitness advantages and were selectively favored regardless of whether the antivirulence compound was present or not. In four cases (ciprofloxacin-gallium, meropenem-gallium, tobramycin-gallium, colistin-furanone), the addition of antivirulence compounds gradually reduced the relative fitness of the AtbR clones, whereby in two cases (meropenem-gallium, tobramycin-gallium) the selective advantage of AtbR clones was completely abrogated. Finally, in one case (tobramycin-furanon) selection for AtbR clones was even reversed and AtbR clones lost the competition.

### Drug synergy does not predict selection against antibiotic resistance

We examined whether drug interactions, ranging from antagonism to synergy for both AtbR clones and the wildtype (Fig 5) correlate with their relative fitness in competition under combination treatments. However, we found no support for such associations (S6 Fig, ANOVA, AtbR: F_1,65_ = 0.88, p = 0.353; WT: F_1,65_ = 1.85, p = 0.179, but instead observed that variation in fitness patterns was explained by specific drug combinations (antivirulence-antibiotic interaction, AtbR: F_3,65_ = 37.45, p < 0.001, WT: F_3,65_ = 14.50, p < 0.001).

### Genetic basis of experimentally evolved antibiotic resistance

The whole-genome sequencing of the experimentally evolved AtbR clones revealed a small number of SNPs and INDELs, which are known to be associated with resistance to the respective antibiotics (Table 1). The AtbR clone resistant to ciprofloxacin had mutations in *gyrB*, a gene encoding the DNA gyrase subunit B, the direct target of the antibiotic [40]. In addition, we identified an 18-bp deletion in the *mexR* gene, encoding a multidrug efflux pump repressor [41]. The two AtbR clones resistant to colistin had different mutations in the same target gene *phoQ* (a non-synonymous SNP in one clone versus a 1-bp insertion in addition to a non-synonymous SNP in the other clone). PhoQ is a regulator of the LPS modification operon and mutations in this gene represent the first step in the development of high-level colistin resistance [42]. One AtbR clone resistant to meropenem had a non-synonymous SNP in the coding sequence of *mpl.* This gene encodes a murein tripeptide ligase, which contributes to the overexpression of the beta-lactamase precursor gene *ampC* [43]. The other AtbR clone resistant to meropenem had mutations in three different genes, which can all be linked to antibiotic resistance mechanisms: we found (i) one non-synonymous SNP in *parR*, which encodes a two component response regulator involved in several resistance mechanisms, including drug efflux, porin loss and LPS modification [44]; (ii) 7 mutations in the *PA1874* gene, which encodes an efflux pump [45]; (iii) one non-synonymous SNP in *nalD*, encoding the transcriptional regulator NalD, which regulates the expression of drug-efflux systems [46,47]. Both AtbR clones resistant to tobramycin had non-synonymous SNPs in *fusA1*. This gene encodes the elongation factor G, a key component of the translational machinery. Although aminoglycosides do not directly bind to the elongation factor G and the complete resistance mechanisms is still unknown, mutations in *fusA1* are associated with high resistance to tobramycin and are often found in clinical isolates [48,49].

**Table 1.**
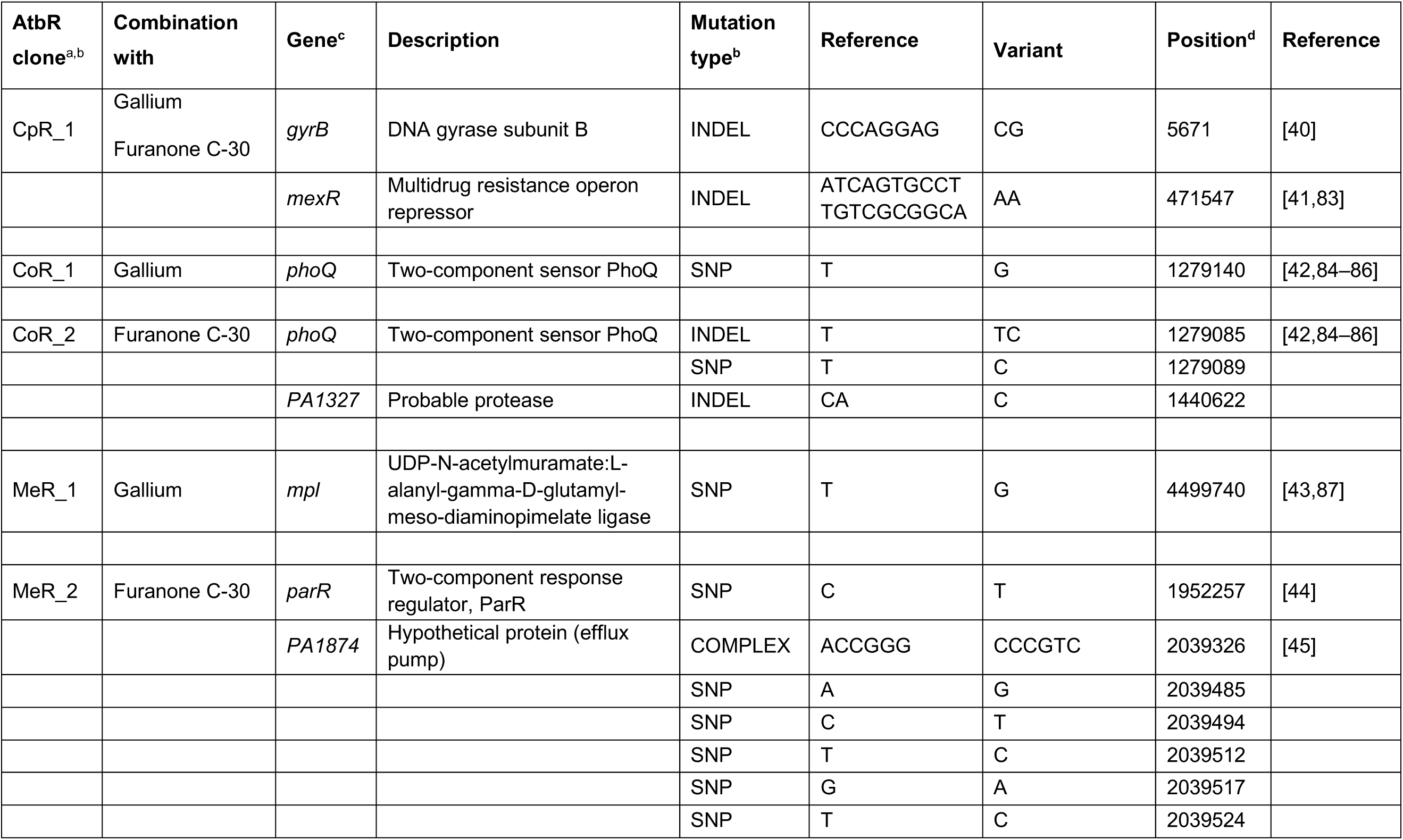
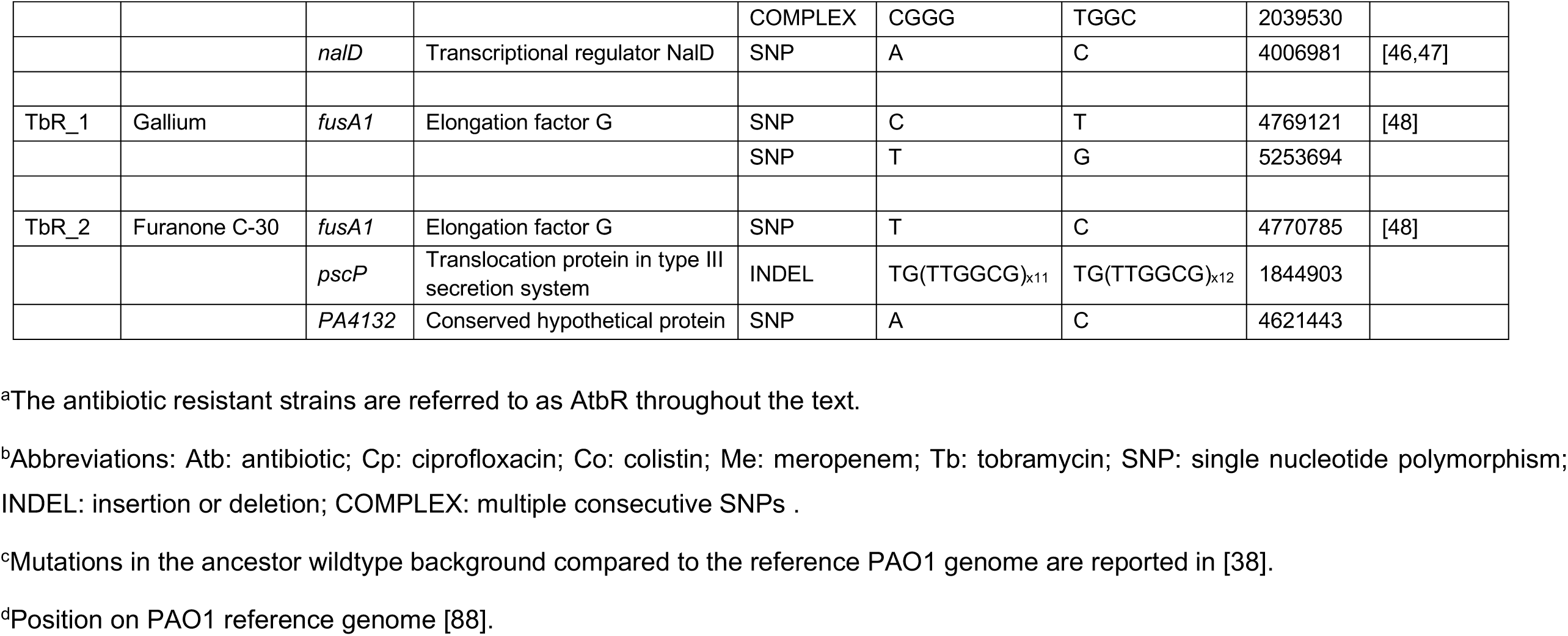
List of mutations in the antibiotic resistant clones.

## Discussion

In this study, we systematically explored the effects of combining antibiotics with antivirulence compounds as a potentially promising strategy to fight susceptible and antibiotic resistant opportunistic human pathogens. Specifically, we combined four different antibiotics (ciprofloxacin, colistin, meropenem, tobramycin) with two antivirulence compounds (gallium targeting siderophore-mediated iron uptake and furanone C-30 targeting the quorum sensing communication system) in 9×9 drug interaction matrices against the bacterium *P. aeruginosa* as a model pathogen. Our heat maps reveal drug-combination specific interaction patterns. While colistin and tobramycin primarily interacted synergistically with the antivirulence compounds, independent and antagonistic interactions occurred for ciprofloxacin and meropenem in combination with the antivirulence compounds (Fig 3+4). We then used antivirulence compounds as adjuvants and observed that they can restore growth inhibition of antibiotic resistant clones in six out of eight cases (Fig 5). Finally, we performed competition assays between antibiotic resistant and susceptible strains under single and combinatorial drug treatments and found that antivirulence compounds can reduce (two cases), abrogate (two cases) or even reverse (one case) selection for antibiotic resistance (Fig 6).

Our results identify antibiotic-antivirulence combinations as a potentially powerful tool to efficiently treat infections of troublesome nosocomial pathogens such as *P. aeruginosa*. From the eight combinations analyzed, tobramycin-antivirulence combinations emerged as the top candidate treatments because: (i) drugs interacted synergistically both with regard to growth and virulence factor inhibition; (ii) the antivirulence compounds restored growth inhibition of antibiotic resistant clones, and even re-potentiated tobramycin in the case of furanone; and (iii) antivirulence compounds either abrogated (for gallium) or even reversed (for furanone) selection for tobramycin resistance, due to collateral sensitivity in the latter case. Meropenem-gallium emerged as an additional promising combination, as this combination also restored growth inhibition of the antibiotic resistant clone and abrogated its selective advantage, despite this drug combination showing slight but significant antagonistic effects.

Drug synergy is desirable from a clinical perspective because it allows to use lower drug concentrations, thereby minimizing side effects while maintaining treatment efficacy [50,51]. In this context, a number of studies have examined combinations of antibiotics and antivirulence compounds targeting various virulence factors including quorum sensing, iron uptake, and biofilm formation in *P. aeruginosa* [12,13,15–18,23,52,53]. While some of these studies have used a few specific concentration combinations to qualitatively assess synergy, we here present comprehensive quantitative interaction maps for these two classes of drugs. A key insight of these interaction maps is that specific drug combinations cannot simply be classified as either synergistic or antagonistic. Instead, drug interactions are concentration dependent with most parts of the interaction maps being characterized by independent effects interspersed with hotspots of synergy or antagonism (Fig 4, S2 Fig). The strongest effects of synergy and antagonism are often observed at intermediate drug concentrations, which is, in the case of synergy, ideal for developing combinatorial therapies that maximize treatment efficacy while minimizing toxicity for the patient. A crucial next step would be to test whether the same type of drug interactions can be recovered in animal host models.

While drug antagonism is considered undesirable from a clinical perspective, work on antibiotic combination therapies has revealed that antagonistic interactions can inhibit the spread of antibiotic resistance [54–56]. The reason behind this phenomenon is that when two drugs antagonize each other, becoming resistant to one drug will remove the antagonistic effect on the second drug, such that the combination treatment will be more effective against the resistant clones [54]. We suspected that such effects might also occur for antagonistic antibiotic-antivirulence treatments, and indeed the meropenem-gallium combination discussed above matches this pattern. However, when comparing across all the eight combinations, we found no evidence that the selection for or against antibiotic resistant clones correlates with the type of drug interaction (S6 Fig). A possible explanation for the overall absence of an association is that the antagonism between antibiotics and antivirulence compounds was quite moderate. In contrast, previous work used an extreme case of antagonism, where the effect of one drug was almost completely suppressed in the presence of the second drug [54,56].

We propose that it is rather the underlying molecular mechanism and not the direction of drug interaction that determines whether selection for antibiotic resistance is compromised or maintained. For instance, any resistance mechanism that reduces antibiotic entry or increases its efflux could conceivably confer cross-resistance to antivirulence compounds, which should in turn maintain or even potentiate and not reverse selection for antibiotic resistance. This phenomenon could explain the patterns observed for furanone in combination with ciprofloxacin and meropenem, where in both cases our sequencing analysis revealed mutations in genes regulating efflux pumps (Table 1). Since furanone needs to enter the cells to become active, these mutations, known to confer resistance to antibiotics [41,42,45], likely also confer resistance to furanone [57]. In contrast, efflux pumps upregulation cannot work as a cross-resistance mechanism against gallium, which binds to secreted pyoverdine and thus acts outside the cell [38].

Alternatively, competitive interactions between resistant and sensitive pathogens over common resources could compromise the spread of drug resistance, as shown for malaria parasites [58]. In our case, it is plausible to assume that antibiotic resistant clones are healthier than susceptible cells and might therefore produce higher amounts of pyoverdine and proteases under antivirulence treatment. Since these virulence factors are secreted and shared between cells, antibiotic resistant clones take on the role of cooperators: they produce costly virulence factors that are then shared with and exploited by the susceptible cells [27,59,60]. This scenario could apply to tobramycin-gallium/furanone combinations, where resistant clones had mutations in *fusA1* known to be associated with the restoration of protein synthesis [48]. Similar social effects could explain selection abrogation in the case of meropenem-gallium combination. Here, the meropenem resistant clone has a mutation in *mpl*, which can trigger the overexpression of the β-lactamase *ampC* resistance mechanism [43]. Since β-lactamase enzyme secretion and extracellular antibiotic degradation is itself a cooperative behaviour [61], it could together with the virulence factor sharing described above compromise the spread of the resistant clone. Clearly, all these explanations remain speculative and further studies are required to understand the molecular and evolutionary basis of abrogated and reversed selection for resistance.

In summary, drug combination therapies are gaining increased attention as more sustainable strategies to treat infections, limiting the spread of antibiotic resistance [62–65]. They are already applied to a number of diseases, including cancer [66], HIV [67] and tuberculosis infections [68]. Here we probed the efficacy and evolutionary robustness of antibiotics combined with anti-virulence compounds. This is an interesting combination because antibiotic treatments alone face the problem of rapid resistance evolution, whereas antivirulence drugs are evolutionarily more robust but can only disarm and not eliminate pathogens. Combinatorial treatments seem to bring the strengths of the two approaches together: efficient removal of bacteria by the antibiotics combined with disarming and increased evolutionary robustness of the antivirulence compounds. While our findings are promising and could set the stage for a novel class of combinatorial treatments, there are still many steps to take to bring our approach to the clinics. First, it would be important to quantify the rate of resistance evolution directly under the combinatorial treatments to test whether drug combination itself slows down resistance evolution [62]. The level of evolutionary robustness of the antivirulence compound would be of particular importance here, as it is known to vary across compounds [27,38]. For example, previous studies showed that resistance to furanone can arise relatively easily [57] while gallium seems to be more evolutionarily robust [38]. Second, the various antibiotic-antivirulence combinations must be tested in relevant animal host models, as host conditions including the increased spatial structure inside the body can affect the competitive dynamics between strains, potentially influencing the outcome of the therapy [69,70]. Relevant in this context would also be tests that closely examine the toxicity of the antivirulence compounds for mammalian cells, an aspect that has received little attention so far [17,71]. Third, our findings suggest that the beneficial effects of combination therapy depend on the specific antibiotic resistance mechanism involved. This hypothesis should be tested in more detail by using sets of mutants that are resistant to the same antibiotic but through different mechanisms. Finally, the observed patterns of drug synergy and reversed selection for resistance are concentration dependent (Fig 4-5). Thus, detailed research on drug delivery and the pharmacodynamics and pharmacokinetics of combination therapies would be required [56], especially to determine the drug interaction patterns within patients.

## Materials and methods

### Bacterial strains

For all our experiments, we used *P. aeruginosa* PAO1 (ATCC 15692). In addition to the wildtype PAO1, we further used two isogenic variants tagged with either a constitutively expressed GFP or mCherry fluorescent protein. Both fluorescently tagged strains were directly obtained from the wildtype PAO1 using the miniTn7-system to chromosomally integrate a single stable copy of the marker gene, under the strong constitutive Ptac promoter, at the *att*Tn7 site [72]. The gentamycin resistance cassette, required to select for transformed clones, was subsequently removed using the pFLP2-encoded recombinase [72]. Antibiotic resistant clones used for competition assays were generated through experimental evolution and are listed in Table 1 together with their respective mutations.

### Media and growth conditions

For all experiments, overnight cultures were grown in 8 ml Lysogeny broth (LB, Sigma Aldrich, Switzerland) in 50 ml Falcon tubes, incubated at 37°C, 220 rpm for 18 hours. We washed overnight cultures with 0.8% NaCl solution and adjusted them to OD_600_ = 1 (optical density at 600 nm). Bacteria were further diluted to a final starting OD_600_ = 10^−3^ for all experiments. We used two different media, where the targeted virulence factors (pyoverdine or protease) are important. For pyoverdine, we used iron-limited CAA (CAA+Tf) [0.5% casamino acids, 5 mM K_2_HPO_4_·3H_2_O, 1 mM MgSO_4_·7H_2_O], buffered at neutral pH with 25 mM HEPES buffer and supplemented with 100 µg/ml human apo-transferrin to chelate iron and 20 mM NaHCO_3_ as a co-factor. As an iron-rich control medium, we used CAA supplemented with 25 mM HEPES and 20 μM FeCl_3_, but without apo-transferrin and 20 mM NaHCO_3_ to create conditions that do not require pyoverdine for growth [36].

For QS-regulated proteases, we used casein medium (CAS) [0.5% casein, 5 mM K_2_HPO_4_·3H_2_O, 1 mM MgSO_4_·7H_2_O], supplemented with 25 mM HEPES buffer and 0.05% CAA. In this medium, proteases are required to digest the casein. A small amount of CAA was added to allow cultures to have a growth kick start prior to protease secretion [73]. As a control, we used CAA supplemented with 25 mM HEPES buffer, a medium in which proteases are not required. All chemicals were purchased from Sigma Aldrich, Switzerland. The CAS medium is intrinsically turbid due to the poor solubility of casein, which can interfere with the growth kinetics measured via optical density (S7A Fig). To solve this issue, we used mCherry fluorescence intensity as a reliable proxy for growth in CAS (S7B-C Fig).

### Single drug growth and virulence factor inhibition curves

To determine the activity range of each antibiotic (ciprofloxacin, colistin, meropenem, tobramycin) and antivirulence drug (gallium as GaNO_3_ and furanone C-30), we subjected PAO1 bacterial cultures to two different 7-step serial dilutions for each antibacterial. Ciprofloxacin: 0-4 μg/ml; colistin: 0-0.4 μg/ml in CAA+Tf and 0-20 μg/ml in CAS; meropenem: 0-14 μg/ml; tobramycin: 0-8 μg/ml; gallium: 0-200 μM; furanone C-30: 0-390 μM. All antibacterials were purchased from Sigma Aldrich, Switzerland. Overnight cultures were prepared and diluted as explained above and then added into 200 μl of media on 96-well plates with six replicates for each drug concentration. Plates were incubated statically at 37°C and growth was measured either as OD_600_ (in CAA+Tf) or mCherry fluorescence (excitation 582 nm, emission 620 nm in CAS) after 48 hours using a Tecan Infinite M-200 plate reader (Tecan Group Ltd., Switzerland). Control experiments (S8 Fig) confirmed that endpoint OD_600_ or mCherry measurements showed strong linear correlations (0.858 < R^2^ < 0.987) with the growth integral (area under the growth curve), which is a good descriptor of the overall inhibitory effects covering the entire growth period [74].

At this time point, we further quantified pyoverdine production through its natural fluorescence (excitation 400 nm, emission 460 nm, a readout that scales linearly with pyoverdine concentration in the medium) and protease production in the cell-free supernatant using the protease azocasein assay (adapted from [38] shortening incubation time to 30 min). The two metals, gallium and bromine (in Furanone C-30), alter the fluorescence levels of pyoverdine and mCherry in a concentration dependent manner. To account for this effect, we established calibration curves and corrected all fluorescence measures accordingly (as described in S9 Fig).

### Antibiotic-antivirulence combination assays

From the single drug dose-response curves, we chose for each drug nine concentrations (including no drugs) to cover the entire activity range in each medium including no, intermediate and strong inhibitory growth effects on PAO1. (S2 Table). We then combined these drug concentrations in a 9×9 matrix for each of the eight antibiotic-antivirulence pairs, and repeated the growth experiment for all combinations in six-fold replication, exactly as described above. After 48 hours of incubation, we measured growth and virulence factor production following the protocols described above.

### Synergy degree of drug-combinations

We used the Bliss independence model to calculate the degree of synergy (*S*), for both growth and virulence factor inhibition, for each of the antibiotic-antivirulence combinations [75–77]. The Bliss model has been suggested as the best model of choice for drugs with different modes of actions and targets [76], as it is the case for antibiotics and antivirulence compounds. We used the formula *S* = *f*_*X*_,_0_. *f* _0_,_*Y*_ − *f* _*X*_,_*Y*_, where *f*_*X,0*_ is the growth (or virulence factor production) level measured under antibiotic exposure at concentration X; *f*_*0,Y*_ is the growth (or virulence factor production) level measured under antivirulence exposure at concentration Y; and *f*_*X,Y*_ is the growth (or virulence factor production) level measured under the combinatorial treatment at concentrations X and Y. If *S* = 0 then the two drugs act independently. Conversely, *S* < 0 indicates antagonistic drug interactions, while S > 0 indicates synergy.

### Experimental evolution under antibiotic treatment

To select for antibiotic resistant clones, we exposed overnight cultures of PAO1 wildtype (initial OD_600_ = 10^−4^) to each of the four antibiotics in LB medium (antibiotic concentrations, ciprofloxacin: 0.15 μg/ml; colistin: 0.5 μg/ml; meropenem: 0.8 μg/ml; tobramycin: 1 μg/ml) in six-fold replication. These antibiotic concentrations initially caused a 70-90% reduction in PAO1 growth compared to untreated cultures, conditions that imposed strong selection for the evolution of resistance. The evolution experiment ran for seven days, whereby we diluted bacterial cultures and transferred them to fresh medium with the respective treatment with a dilution factor of 10^−4^, every 24 hours. At the end of each growth cycle, we measured growth (OD_600_) of the evolving lineages using a SpectraMax® Plus 384 plate reader (Molecular Devices, Switzerland).

### Phenotypic and genetic characterization of resistance

Following experimental evolution, we screened the evolved lines for the presence of antibiotic resistant clones. For each antibiotic we plated four evolved lines on LB plates and isolated single clones, which we then exposed in liquid culture to the antibiotic concentration they experienced during experimental evolution. Among those that showed growth restoration (compared to the untreated wildtype), we picked two random clones originating from different lineages per antibiotic for further analysis. We had to adjust our sampling design in two cases. First, only one population survived our ciprofloxacin treatment and thus only one resistant clone could be picked for this antibiotic. Second, clones evolved under colistin treatment grew very poorly in CAS medium and therefore we included an experimentally evolved colistin resistant clone from a previous study, which did not show compromised growth in CAS (see [38] for a description on the experimental evolution). Altogether, we had seven clones (one clone per antibiotic was allocated to one of the two media, except for ciprofloxacin). For all of these clones, we re-established the drug-response curves in either CAA+Tf or CAS and quantified the IC50 values (S4 Fig). For all cases, the IC50 of the AtbR clones was significantly higher than the ones of the antibiotic-sensitive wildtype (−187.30 ≤ *t* ≤ - 3.10, p < 0.01 for all AtbR clones; S4 Fig). Furthermore, we examined whether resistance to antibiotics can lead to collateral sensitivity or cross-resistance to antivirulence compounds, and that is why also established the dose-response curves for all the seven AtbR clones for the respective antivirulence compounds. As before, we exposed the clones and the wildtype PAO1 to a range of gallium (0-50 μM) or furanone (0-390 μM) concentrations in CAA+Tf or CAS, respectively, and compared their IC50 values.

We further isolated the genomic DNA of the selected evolved antibiotic resistant clones and sequenced their genomes. We used the GenElute Bacterial Genomic DNA kit (Sigma Aldrich) for DNA isolation. DNA concentrations were assayed using the Quantifluor dsDNA sample kit (Promega, Switzerland). Samples were sent to the Functional Genomics Center Zurich for library preparation (TruSeq DNA Nano) and sequencing on the Illumina MiSeq platform with v2 reagents and pair-end 150 bp reads. In a first step, we mapped the sequences of our ancestral wildtype PAO1 strain (ENA accession number: ERS1983671) to the *Pseudomonas aeruginosa* PAO1 reference genome (NCBI accession number: NC_002516) with snippy (https://github.com/tseemann/snippy) to obtain a list with variants that were already present at the beginning of the experiment. Next, we quality filtered the reads of the evolved clones with trimmomatic [78], mapped them to the reference genome and called variants using snippy. Detected variants were quality filtered and variants present in the ancestor strain were excluded from the dataset using vcftools [79]. The mapping files generated in this study are deposited in the European Nucleotide Archive (ENA) under the study accession number PRJEB32766.

### Monoculture and competition experiments between sensitive and resistant clones

To examine the effects of combination treatments on the growth and the relative fitness of antibiotic resistant clones, we subjected the sensitive wildtype PAO1 (tagged with GFP) and the experimentally evolved antibiotic resistant clones (Table 1), alone or in competition, to six different conditions: (i) no drug treatment; (ii) antibiotics alone; (iii-iv) two concentrations of antivirulence compounds alone and (v-vi) the same two concentrations of antivirulence compounds combined with antibiotics. Antibiotic concentrations are listed in the S2 Table, while antivirulence concentrations were as follows, gallium: 1.56 μM (low), 6.25 μM (intermediate); furanone: 6.3 μM (low), 22.8 μM (intermediate). Bacterial overnight cultures were prepared and diluted as described above. Competitions were initiated with a mixture of 90% sensitive wildtype cells and 10% resistant clones to mimic a situation where resistance is still relatively rare. Mixes alongside with monocultures of all strains were inoculated in either 200 μl of CAA+Tf or CAS under all the six treatment regimes. We used flow cytometry to assess strain frequency prior and after a 24 hours competition period at 37°C static (S10 Fig). Specifically, bacterial cultures were diluted in 1X phosphate buffer saline (PBS, Gibco, ThermoFisher, Switzerland) and frequencies were measured with a LSRII Fortessa cell analyzer (BD Bioscience, Switzerland. GFP channel, laser: 488 nm, mirror: 505LP, filter: 530/30; side and forward scatter: 200 V threshold; events recorded with CS&T settings) at the Cytometry Facility of the University of Zurich. We recorded 50’000 events before competitions and used a high-throughput sampler device (BD Bioscience) to record all events in a 5 μl-volume after competition. Since antibacterials can kill and thereby quench the GFP signal in tagged cells, we quantified dead cells using the propidium iodide (PI) stain (2 μl of 0.5 mg/ml solution) with flow cytometry (for PI fluorescence: laser: 561 nm, mirror: 600LP, filter: 610/20).

We used the software FlowJo (BD Bioscience) to analyse data from flow cytometry experiments. We followed a three-step gating strategy: (i) we separated bacterial cells from media and noise background by using forward and side scatter values as a proxy for particle size; (ii) within this gate, we then distinguished live from dead cells based on the PI staining; (iii) finally, we separated live cells into either GFP positive and negative populations. Fluorescence thresholds were set using appropriate control samples: isopropanol-killed cells for PI positive staining and untagged-PAO1 cells for GFP-negative fluorescence. We then calculated the relative fitness of the antibiotic resistant clone as ln(v) = ln{[a_24_×(1−a_0_)]/[a_0_×(1−a_24_)]}, where a_0_ and a_24_ are the frequencies of the resistant clone at the beginning and at the end of the competition, respectively [80]. Values of ln(v) < 0 or ln(v) > 0 indicate whether the frequency of antibiotic resistant clones decreased or increased relative to the sensitive PAO1-GFP strain. To check for fitness effects caused by the fluorescent tag, we included a control competition, where we mixed PAO1-GFP with the untagged PAO1 in a 9:1 ratio for all treatment conditions. We noted that high drug concentrations significantly curbed bacterial growth, which reduced the number of events that could be measured with flow cytometry. This growth reduction increased noise relative to the signal, leading to an overestimation of the GFP-negative population in the mix. To correct for this artefact, we established calibration curves for each individual experimental replicate for how the relative fitness of PAO1-untagged varies as a function of cell density in control competitions with PAO1-GFP. Coefficients of the asymptotic functions used for the correction are available together with the raw dataset.

### Statistical analysis

All statistical analyses were performed with RStudio v. 3.3.0 [81]. We fitted individual dose response curves with either log-logistic or Weibull functions, and estimated and compared IC50 values using the drc package [82], while dose response curves under combination treatment were fitted using spline functions. We used Pearson correlation coefficients to test for significant associations between the degree of synergy in growth and virulence factor inhibition. We used one-sample t-tests to compare the degree of synergy of each combination of concentrations represented in Fig 4 to zero. We used Welch’s two-sample t-test to compare growth between the sensitive wildtype PAO1 and the resistant clones under antibiotic treatment. To compare the relative fitness of resistant clones to the reference zero line, we used one-sample t-tests. Finally, we used analysis of variance (ANOVA) to test whether the addition of antivirulence compounds to antibiotics affected the relative fitness of AtbR clones, and whether the outcome of the competition experiment is associated with the degree of synergy of the drug combinations. Where necessary, p-values were adjusted for multiple comparisons using the false discovery rate method.

## Acknowledgments

We thank Roland Regös and Désirée Bäder for advice on the Bliss model; Alex Hall, Roland Regös and Frank Schreiber for comments on the manuscripts; David Wilson for experimental support; Selina Niggli, Priyanikha Jayakumar and the Flow Cytometry Facility (University of Zurich) for support with flow cytometry experiments; and the Functional Genomics Center Zurich for technical support with the strain sequencing.

