## Supporting Information for "Combining antibiotics with antivirulence compounds can have synergistic effects and reverse selection for antibiotic resistance in *Pseudomonas aeruginosa*"

**Short title**: Antibiotic-antivirulence combination therapy against bacterial pathogens

**Corresponding author**: Rolf Kümmerli, Department of Quantitative Biomedicine, University of Zurich, Winterthurerstrasse 190, 8057 Zurich, Switzerland.

This document contains:

- 10 supplementary figures
- 2 supplemetary tables

### **Supplementary Figures**

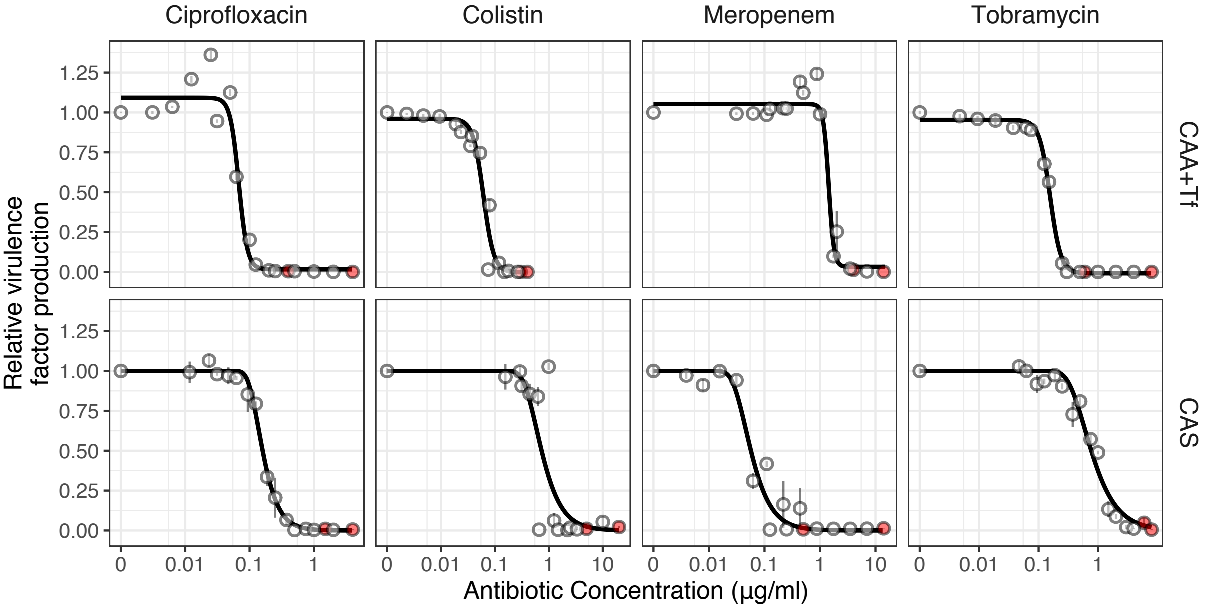


**S1 Figure. Effect of antibiotics on virulence factor production in *P. aeruginosa* PAO1 populations.** We exposed PAO1 to all four antibiotics in two the experimental media: CAA+Tf (iron-limited casamino acid medium with transferrin) and CAS (casein medium). After 48 h exposure, we measured virulence factor production: pyoverdine in CAA+Tf and proteases in CAS. The inhibition of virulence factors followed the same pattern as for growth inhibition, except for ciprofloxacin and meropenem, where pyoverdine production slightly increased at intermediate antibiotic concentrations and only dropped at higher antibiotic levels. Dots show means ± standard error across six replicates. All data are scaled relative to the drug-free treatment. Data stem from the same two independent experiments as shown in Figure 1. The red dots indicate the highest concentration used for each experiment, from which 7-serial dilution steps were tested. Curves were fitted with log-logistic functions.

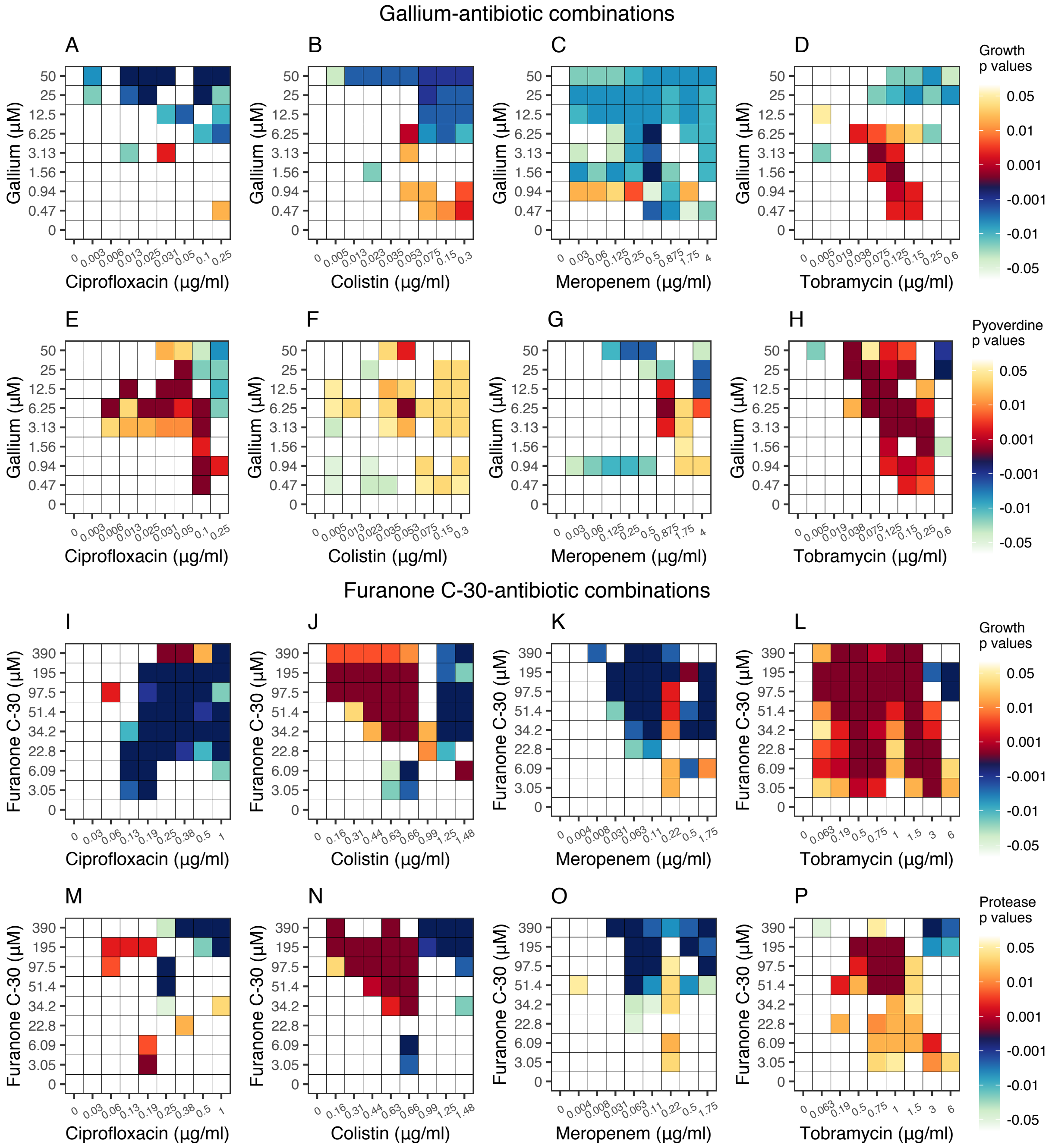


**S2 Figure. Statistical significance maps for antibiotic-antivirulence drug interactions.** For each drug concentration combination, we tested whether the degree of synergy is significantly different from zero (i.e. independent drug interaction). Heatmaps depict p-values ranging from white (no significant drug interaction) to blue (significant antagonism) to red (significant synergy). P-values are shown for gallium-antibiotic combinations (**A-D** for growth; **E-H** for pyoverdine production) and furanone-antibiotic combinations (**I-L** for growth; **M-P** for protease production). To account for multiple comparisons, we corrected the p-values for each drug combination using the "false discovery rate" method.

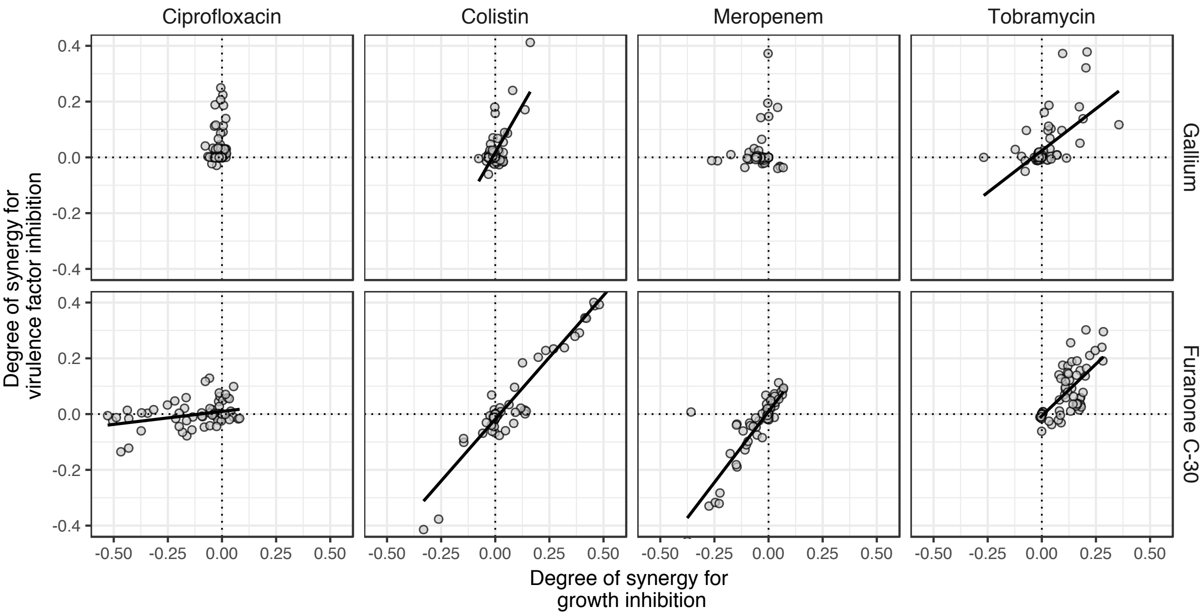


**S3 Figure. Assessing the relationship between the degrees of synergy for growth and virulence factor inhibition.** We found that the degrees of synergy for the two measured traits across the 9x9 antibiotic-antivirulence combination matrix correlated in 6 out of 8 cases (Pearson correlation coefficient: ciprofloxacin-gallium: r = 0.09, $t_{79}$ = 0.85, p = 0.394; colistin-gallium: r = 0.69, $t_{79}$ = 8.51, p < 0.001; meropenem-gallium: r = 0.17, $t_{79}$ = 1.52, p = 0.130; tobramycin-gallium: r = 0.58, $t_{79}$ = 6.39, p < 0.001; ciprofloxacin-furanone: r = 0.34, $t_{79}$ = 3.22, p = 0.002; colistin-furanone: r = 0.96, $t_{79}$ = 32.50, p < 0.001; meropenem-furanone: r = 0.87, $t_{79}$ = 15.48, p < 0.001; tobramycin-furanone: r = 0.75, $t_{79}$ = 10.16, p < 0.001). Solid lines show association trend lines between the two levels of interactions.

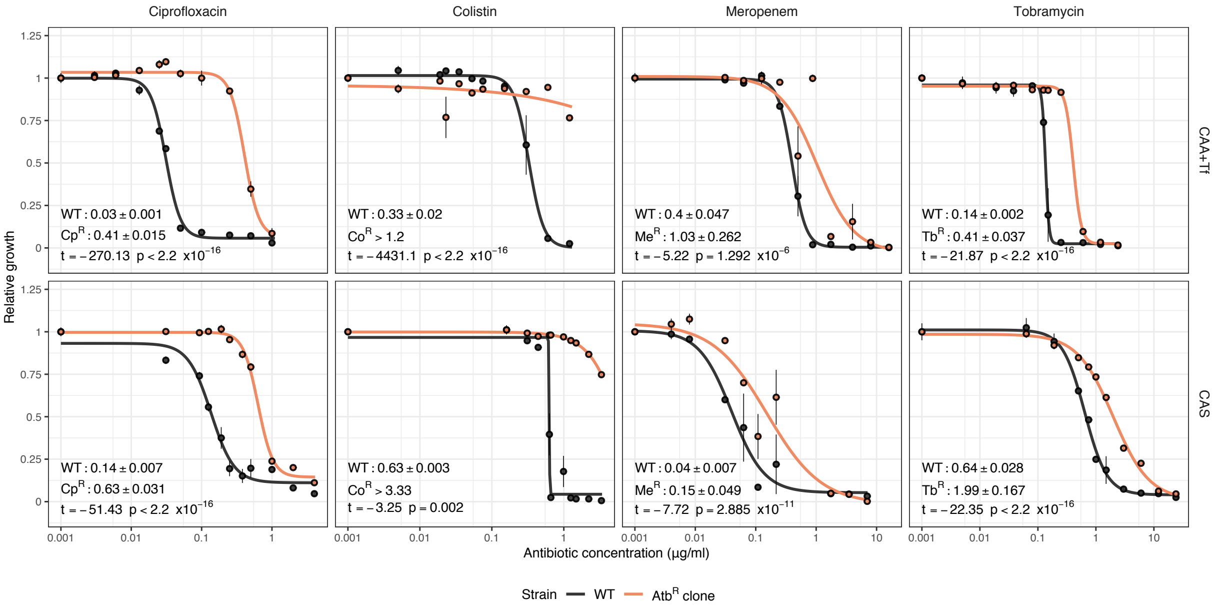


**S4 Figure. Clones evolved under antibiotic treatments show altered dose response curves.** To confirm that the evolved clones are resistant to the antibiotic in the two experimental media (iron limited (CAA+Tf) and casein (CAS) media), we measured for each antibiotic the dose-response curves for the ancestral wildtype and one randomly selected clone either in CAA+Tf or CAS. For each antibiotic and medium, we tested 11 concentrations within these ranges: ciprofloxacin: 0-1 µg/ml (CAA+Tf), 0-4 µg/ml (CAS); colistin: 0-1.2 µg/ml (CAA+Tf), 0-3.33 µg/ml (CAS); meropenem: 0-1.6 µg/ml (CAA+Tf), 0-7 µg/ml (CAS); tobramycin: 0-2.4 µg/ml (CAA+Tf), 0-24 µg/ml (CAS). All evolved clones showed an attenuated dose-response curve, were able to grow at higher drug concentrations than the ancestral wildtype (WT) and had significantly higher half maximal inhibitory concentration (IC50) values. Measurements of $OD_{600}$ were taken after 48 hours incubation time at 37°C under static conditions. Growth values are scaled relative to the untreated control for each strain. Data are shown as means ± standard errors across four replicates. Curves were fitted with four parameters log-logistic functions. In the lower left corner of each panel we show the mean IC50 values of the WT and AtbR clones ± standard errors (µg/ml) and the respective statistical analysis comparing the ratio of the means.

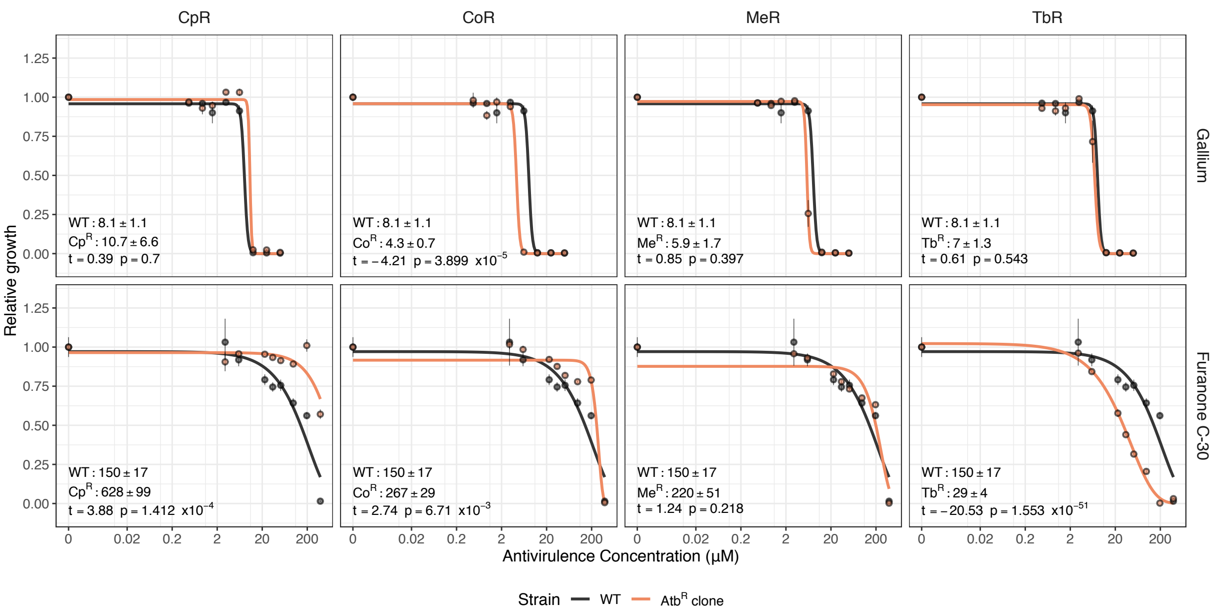


**S5 Figure. Antivirulence dose-response curves for antibiotic resistant clones of *P. aeruginosa* PAO1.** To check whether resistance to antibiotics influenced the susceptibility to antivirulence compounds, we exposed our selected antibiotic resistant (AtbR) clones to a range of concentrations of both gallium (0-50 µM) and furanone (0-390 µM). Under gallium treatment, only the colistin resistant clone showed increased sensitivity to the antivirulence drug. Under furanone treatment, the clones resistant to ciprofloxacin and colistin showed a certain level of cross-resistance to this antivirulence compound, while we found collateral sensitivity between tobramycin and furanone. All values are scaled relative to the untreated control for each strain, and data points show the mean across four replicates. We used either log-logistic functions (in CAA+Tf) or three-parameter Weibull functions (in CAS) to fit the curves and extract mean IC50 values ± standard error, which are reported in the left bottom corner of each panel together with the respective statistical analysis comparing the ratio of the means.

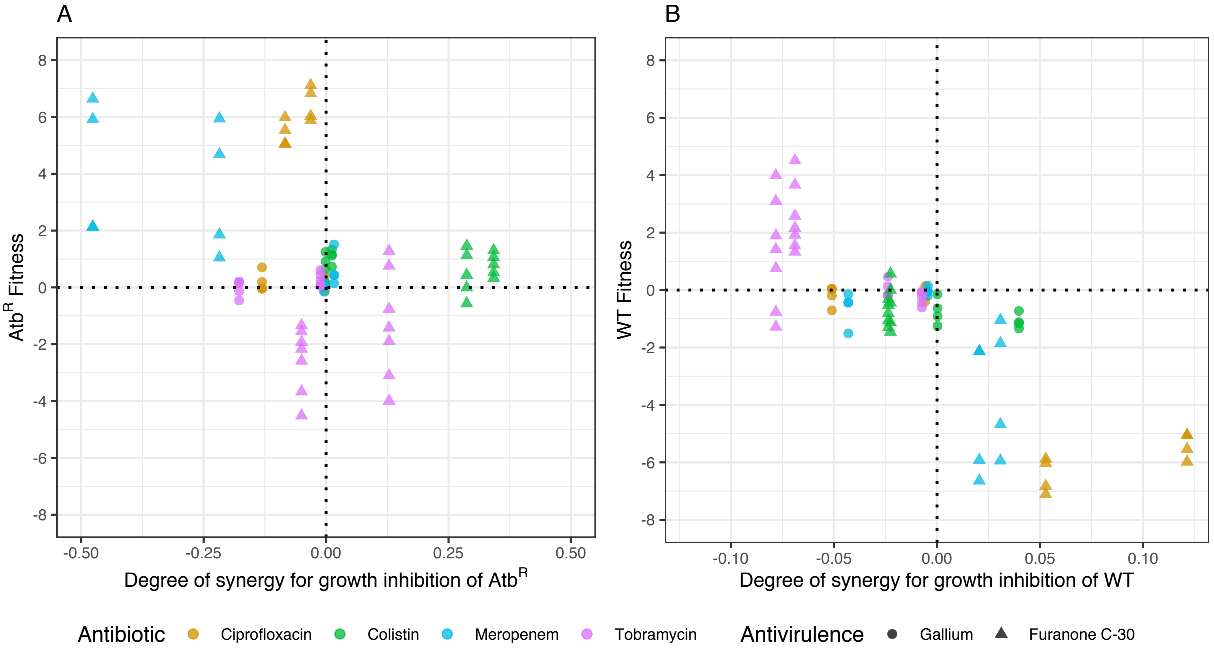


**S6 Figure. Testing for correlations between the degree of synergy for growth inhibition and the outcome of competitions.** We tested whether the degree of synergy for growth inhibition is a predictor of the competition outcome between the antibiotic resistant clones (AtbR) and the susceptible wild type (WT) under combination treatment. We compared the degrees of synergy of each drug combination for the AtbR clones (**A**) or the WT (**B**) to their relative fitness values in competition. Positive or negative y-values indicate that the clones increased or decreased in frequency during the competition, respectively. Positive or negative values on the x-axis indicate synergy or antagonism, respectively. There were no significant associations between the relative fitness and the degree of synergy for growth inhibition neither for the AtbR clones nor for the WT (ANOVA, for AtbR: $F_{1,65} = 0.88$, $p = 0.353$; for WT: $F_{1,65} = 1.85$, $p = 0.179$). Instead, relative fitness was significantly affected by the type of antivirulence drug (ANOVA, for AtbR: $F_{1,65} = 106.36$, $p < 0.001$; for WT: $F_{1,65} = 44.58$, $p < 0.001$) and the specific antibiotic-antivirulence combination applied (ANOVA, for AtbR: $F_{3,65} = 37.45$, $p < 0.001$; for WT: $F_{3,65} = 14.50$, $p < 0.001$).

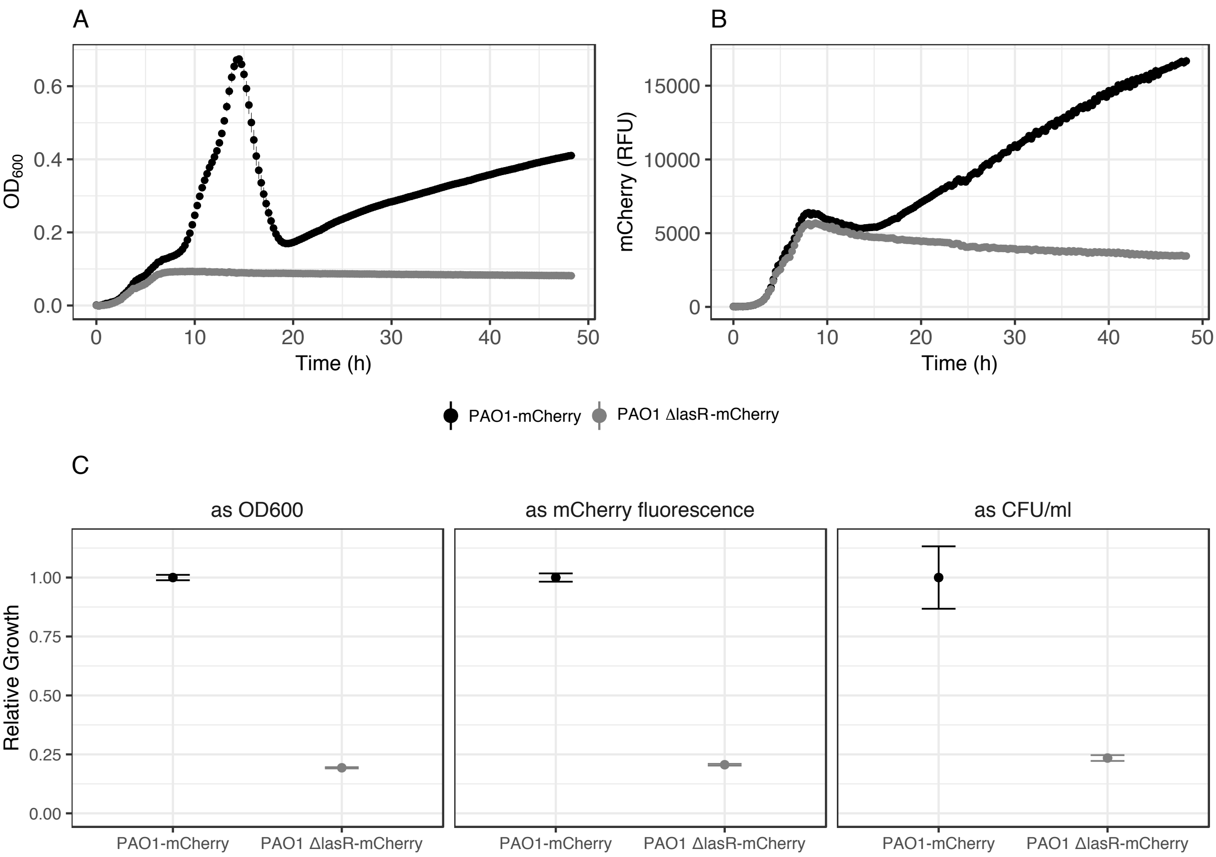


**S7 Figure. Validation of mCherry fluorescence as a proxy for growth measurements.** The CAS (casein) media has a very high turbidity due to the poor solubility of casein, which interferes with optical density at 600 nm ($OD_{600}$), which is typically used as a measure of growth. We therefore used mCherry fluorescence, constitutively expressed from a single-copy chromosomal insertion, as a proxy for bacterial growth in CAS. To validate this method, we grew PAO1-mCherry (able to digest CAS) and PAO1 *ΔlasR*-mCherry (unable to digest CAS) in CAS medium for 48 hours at 37°C in a Tecan plate reader tracking $OD_{600}$ and mCherry fluorescence every 15 minutes. (**A**) Blank corrected $OD_{600}$ trajectories for PAO1-mCherry and PAO1 *ΔlasR*-mCherry. PAO1 *ΔlasR*-mCherry grew poorly but showed a standard sigmoid growth pattern by digesting the supplemented CAA. In stark contrast, the $OD_{600}$ of PAO1-mCherry first increased sharply, then declined dramatically followed by a slow linear increase over time. This trajectory is explained by the simultaneous growth of bacteria (increasing $OD_{600}$) and clearance of the turbidity due to protein digestion (decreasing $OD_{600}$), thus demonstrating that $OD_{600}$ is an unsuitable measure for growth. (**B**) Blank corrected mCherry trajectories for PAO1-mCherry and PAO1 *ΔlasR*-mCherry. As for $OD_{600}$, PAO1 *ΔlasR*-mCherry grew only poorly (according to the mCherry signal) and only within the first 7 hours of the assay, digesting the supplemented CAA. Unlike for $OD_{600}$, the mCherry signal yielded a much more sensible growth trajectory for PAO1-mCherry, characterized by an initial increase (CAA consumption), followed by a lag phase (protease secretion and switch to CAS) and growth resumption (CAS digestion). (**C**) To further validate that mCherry fluorescence is a good proxy for growth in CAS media, we compared the endpoints measurements of mCherry fluorescence with

CFU/ml values, determined by plating the cultures on LB-agar plates. All values are scaled relative to PAO1-mCherry. The two methods yielded similar results and show that the growth of PAO1*ΔlasR*-mCherry is approximately 25% of the one of PAO1-mCherry. Data are shown as means ± standard errors across eight replicates for the growth curves and eight (PAO1-mCherry) or four (PAO1*ΔlasR*-mCherry) replicates for the CFU/ml data.

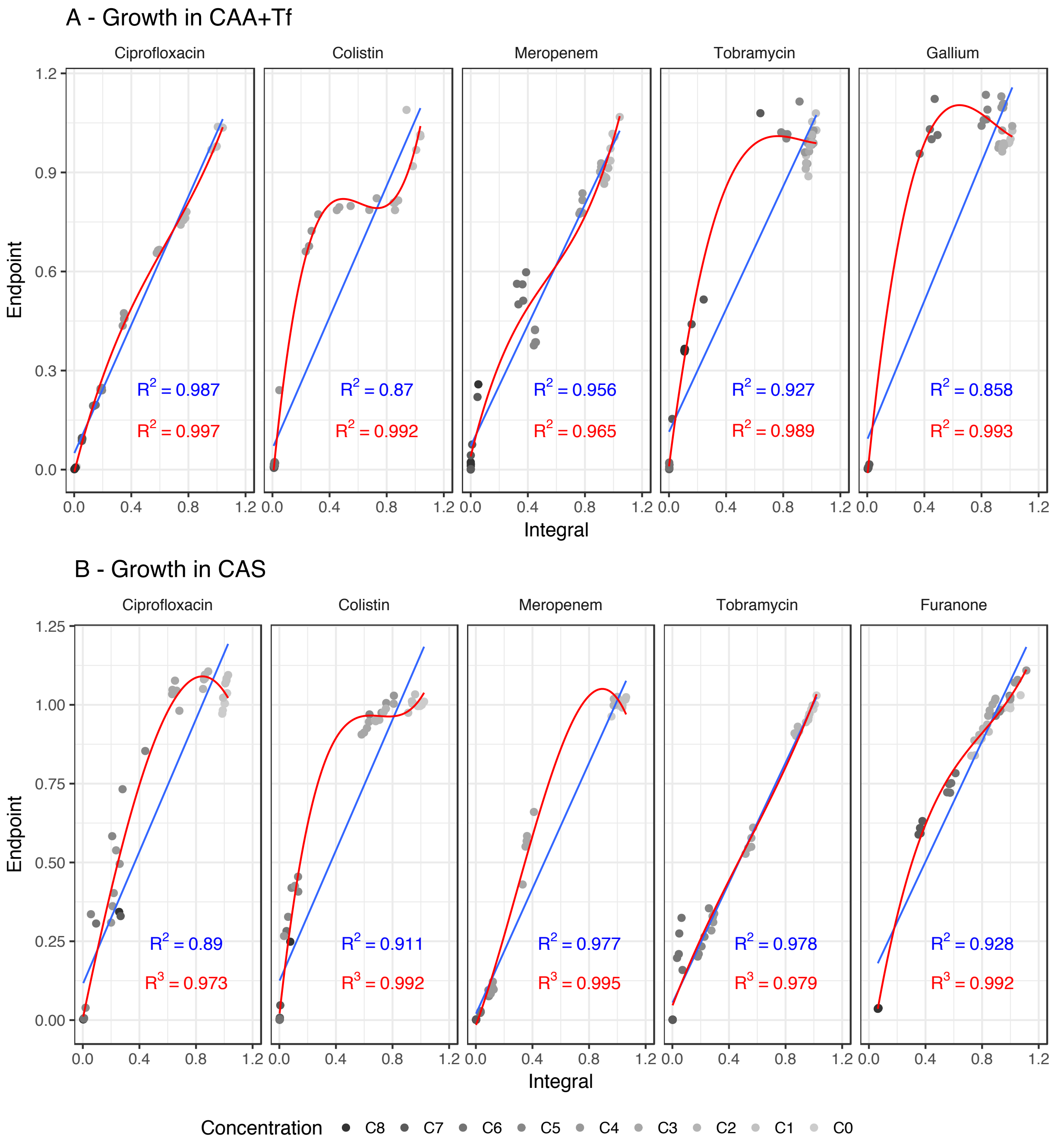


**S8 Figure. Correlation between endpoint measurements and area under the growth curve (integral) under single drug treatments.** To verify that a single $OD_{600}$ or mCherry measurement after growth is a good proxy for growth inhibition, we tested the correlation between the area under the growth curve (integral) and endpoint measurements, under single drug treatments in CAA+Tf (**A**) or CAS (**B**) media. For each antibiotic and antivirulence compound we picked 9 concentrations which cover the entire drug active range, and that were used for the combination assay, shown in Figure 3 and Figure 4. Each concentration was tested in 5-fold replication. Cultures were grown for 48 hours in a Tecan Infinite M-200 plate reader (Tecan Group Ltd., Switzerland) and growth was recorded by reading $OD_{600}$ (in CAA+Tf) or mCherry fluorescence (in CAS) every 15 min, after a short shaking event. Growth trajectories were established with a spline fit and the two parameters (endpoint yield and integral) were extracted using the grofit package in RStudio [88]. In both media, the two growth parameters showed strong linear association patterns (blue lines and $R^2$ values). For several drugs, growth integral measurements were more sensitive to discover growth inhibitions at low drug concentrations (light grey circles) and that is why cubic data fits (red lines and $R^2$ values) often explained an even higher proportion of the variance. Nonetheless, these control analyses show that endpoint growth values are reliable proxies for measuring growth inhibition under drug treatment.

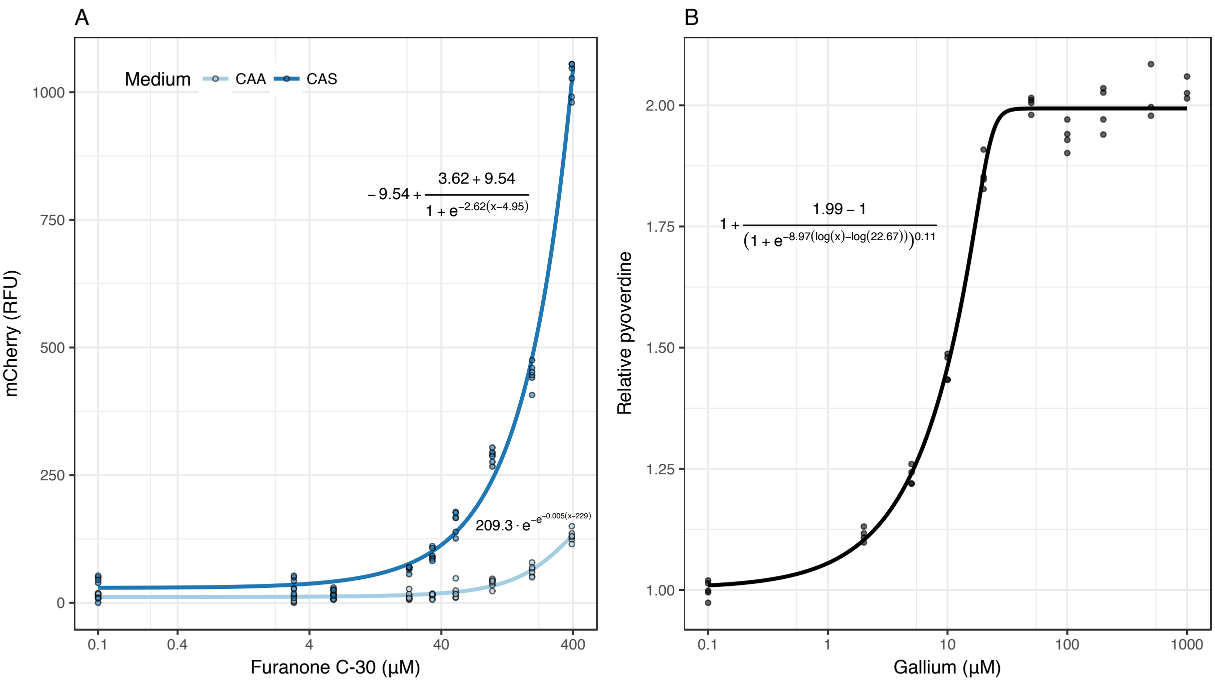


**S9 Figure. Fluorescence correction for mCherry and pyoverdine measurements in the presence of furanone C-30 and gallium**. The two metals, bromine (in furanone C-30) and gallium interfere with the fluorescence measurements of mCherry and pyoverdine in a concentration dependent manner. To account for this bias, we established calibration curves and used them to correct fluorescent values in all experiments. (**A**) Furanone C-30 is autofluorescent in the mCherry channel (excitation 582 nm, emission 620 nm). We quantified the autofluorescence in function of the concentration of furanone both in CAS (casein) and CAA (casamino acids) media. Briefly, we incubated each media supplemented with a range of furanone C-30 concentrations (0-390 µM, as used in Figure 2, in 6-fold replication) for 48 hours under static conditions and then measured mCherry fluorescence. The relationship between concentration and fluorescence was explained by a four parameters logistic function in CAS or by a three parameters Gompertz function in CAA. In all experiments, we used this calibration curve to subtract, for each furanone concentration, the autofluorescence component from the mCherry measurements. (**B**) The fluorescent signal of pyoverdine becomes inflated when gallium binds to the siderophore [26,73]. We used the supplementary data from Ross-Gillespie *et al*. [26] to quantify this bias in fluorescence as a function of gallium concentration. They incubated 200 µM of pyoverdine in iron limited CAA+Tf medium, supplemented with gallium concentrations ranging from 0 to 1 mM and measured pyoverdine-associated fluorescence. When supplemented with more than 50 µM of gallium, pyoverdine showed a nearly 2-fold higher fluorescence signal. This signal bias can be explained by a five parameters log-logistic function. In all our experiments, for each gallium concentration used, we applied correction factors derived from this fitted curve to account for this potential bias.

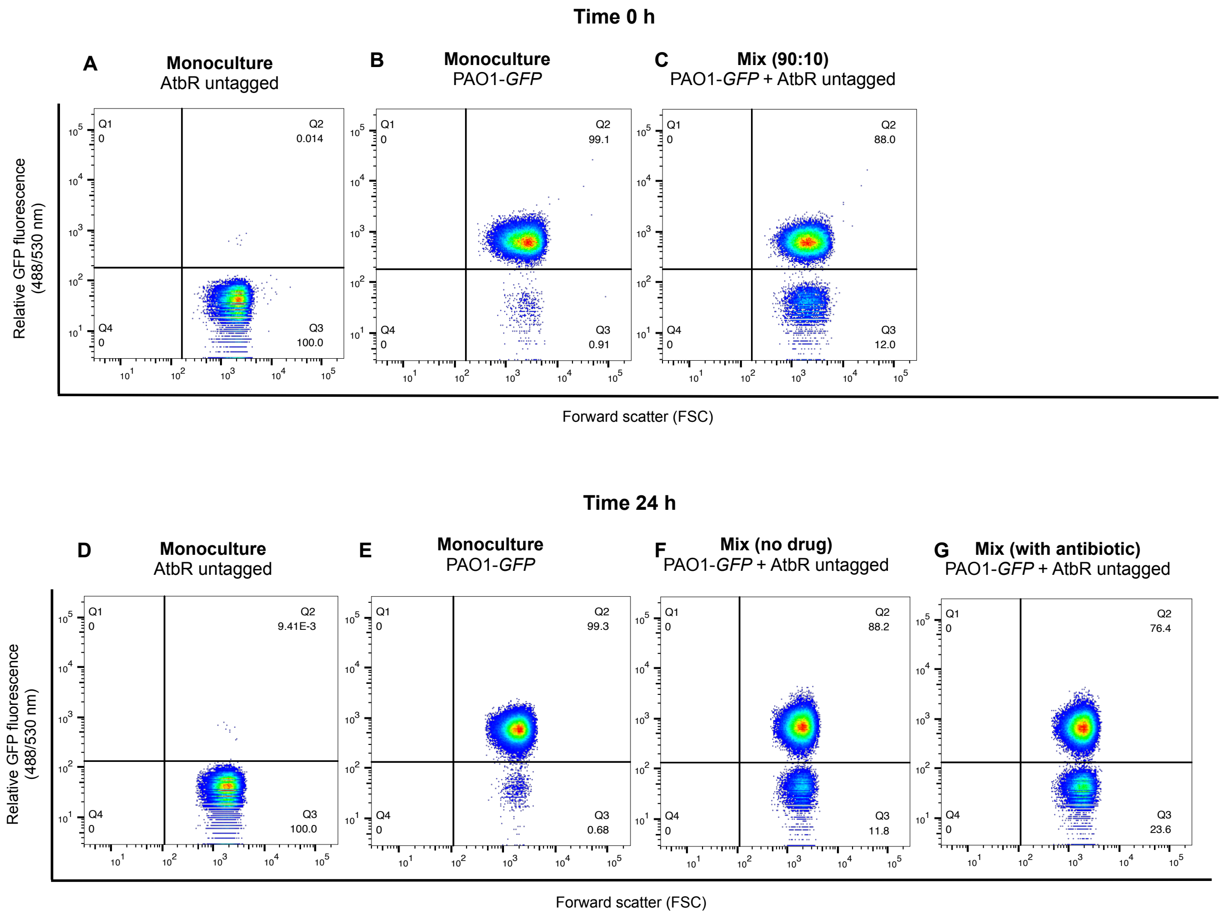


**S10 Figure. Examples of flow-cytometry scatter plots from competition experiments between the sensitive wildtype PAO1 and antibiotic resistant clones.** The wildtype strain PAO1, chromosomally tagged with a constitutively expressed GFP marker, was co-cultured with antibiotic resistant clones (AtbR) in a 90:10 ratio in the presence of five different drug treatments. Mono- and mixed- cultures were measured with the flow cytometer at the beginning (time 0 h) and at the end (time 24 h) of the competition experiments. For data analysis, we plotted the size of the cells (forward scatter, FSC) against the GFP fluorescence to distinguish tagged from untagged cells. The shown plots depict an illustrative example, in which ciprofloxacin was used as an antibiotic. (**A**) A monoculture of the untagged AtbR clone does not show GFP fluorescence. This control allows to quantify the background fluorescence of the cells. (**B**) A monoculture of the tagged PAO1-GFP shows relatively strong GFP fluorescence, with 99.1% of all cells considered as GFP positive. (**C**) In a 90:10 volumetric mix of wildtype and AtbR clones, the cells of the two strains can be unambiguously distinguished and their actual ratio (88:12) can be determined. Frequencies of GFP-positive and GFP-negative cells were then quantified after a 24 hours incubation period at 37°C. (**D**) The monoculture of the untagged AtbR strain shows that cells do not increase their GFP autofluorescence over time and 100% of cells fall into the GFP-negative gate. (**E**) The monoculture of PAO1-GFP shows relatively strong fluorescence also at the end of the competition, with 99.3% of cells being classified as GFP positive. (**F**) The mix of wildtype and AtbR clones, when grown in absence of any drug treatment stays at the initial frequency (88.2:11.8). (**G**) When the mix was grown in the presence of the antibiotic, the fraction of untagged AtbR strain increases to 23.6%, demonstrating their selective advantage

**S1 Table. Statistical analyses (t-test and ANOVA) performed on the data presented in Figure 6**

| Combination | Antibiotic presence | Factor[2] | Single comparisons[1,2] | Degrees of freedom[1,2] | F value[2] | t value[1,2] | p-value[1,2] |
|---|---|---|---|---|---|---|---|
| Ciprofloxacin-Gallium | | | | | | | |
| | - Atb | | 0 vs reference line | 15 | | -5.71 | $4.121 \times 10^{-5}$ |
| | | | Low vs reference line | 7 | | -13.56 | $2.786 \times 10^{-6}$ |
| | | | Intermediate vs reference line | 7 | | 2.51 | 0.040 |
| | + Atb | | 0 vs reference line | 15 | | 7.27 | $2.749 \times 10^{-6-}$ |
| | | | Low vs reference line | 15 | | 2.54 | 0.022 |
| | | | Intermediate vs reference line | 15 | | 2.17 | 0.046 |
| | | Treatment | | 2 | 4.17 | | 0.022 |
| | | Residuals | | 45 | | | |
| | | | 0 vs Low | | | -1.82 | 0.076 |
| | | | 0 vs Intermediate | | | -2.85 | 0.007 |
| Colistin-Gallium | | | | | | | |
| | - Atb | | 0 vs reference line | 15 | | -13.50 | $8.518 \times 10^{-10}$ |
| | | | Low vs reference line | 7 | | 8.86 | $4.718 \times 10^{-5}$ |
| | | | Intermediate vs reference line | 7 | | 3.16 | 0.016 |
| | + Atb | | 0 vs reference line | 15 | | 13.12 | $1.266 \times 10^{-9}$ |
| | | | Low vs reference line | 15 | | 17.15 | $2.901 \times 10^{-11}$ |
| | | | Intermediate vs reference line | 15 | | 7.02 | $4.139 \times 10^{-6}$ |

| | | | | | | | |
|---|---|---|---|---|---|---|---|
| | | Treatment | | 2 | 15.31 | | $8.482 \times 10^{-6}$ |
| | | Residuals | | 45 | | | |
| | | | 0 vs Low | | | 5.49 | $1.760 \times 10^{-6}$ |
| | | | 0 vs Intermediate | | | 2.16 | 0.036 |
| Meropenem-Gallium | | | | | | | |
| | - Atb | | 0 vs reference line | 15 | | -6.20 | $1.691 \times 10^{-5}$ |
| | | | Low vs reference line | 7 | | -1.87 | 0.104 |
| | | | Intermediate vs reference line | 7 | | -4.48 | 0.003 |
| | + Atb | | 0 vs reference line | 15 | | 5.80 | $3.480 \times 10^{-5}$ |
| | | | Low vs reference line | 15 | | 4.65 | $3.140 \times 10^{-4}$ |
| | | | Intermediate vs reference line | 15 | | 0.72 | 0.482 |
| | | Treatment | | 2 | 12.30 | | $5.495 \times 10^{-5}$ |
| | | Residuals | | 45 | | | |
| | | | 0 vs Low | | | -0.87 | 0.390 |
| | | | 0 vs Intermediate | | | -4.66 | $2.810 \times 10^{-5}$ |
| Tobramycin-Gallium | | | | | | | |
| | - Atb | | 0 vs reference line | 19 | | -6.30 | $4.768 \times 10^{-6}$ |
| | | | Low vs reference line | 7 | | -2.49 | 0.042 |
| | | | Intermediate vs reference line | 7 | | -3.90 | 0.006 |
| | + Atb | | 0 vs reference line | 19 | | 8.42 | $7.763 \times 10^{-8}$ |
| | | | Low vs reference line | 19 | | -0.16 | 0.877 |

| | | | | | | | |
|---|---|---|---|---|---|---|---|
| | | | Intermediate vs reference line | 19 | | 5.71 | $1.675 \times 10^{-5}$ |
| | | Treatment | | 2 | 22.99 | | $4.766 \times 10^{-8}$ |
| | | Residuals | | 57 | | | |
| | | | 0 vs Low | | | -6.78 | $7.360 \times 10^{-9}$ |
| | | | 0 vs Intermediate | | | -3.38 | 0.001 |
| Ciprofloxacin-Furanone C30 | | | | | | | |
| | - Atb | | 0 vs reference line | 15 | | -12.56 | $2.312 \times 10^{-9}$ |
| | | | Low vs reference line | 7 | | -10.50 | $1.546 \times 10^{-5}$ |
| | | | Intermediate vs reference line | 7 | | -0.22 | 0.835 |
| | + Atb | | 0 vs reference line | 15 | | 13.41 | $9.315 \times 10^{-10}$ |
| | | | Low vs reference line | 15 | | 35.13 | $8.04 \times 10^{-16}$ |
| | | | Intermediate vs reference line | 15 | | 51.13 | $< 2.2 \times 10^{-16}$ |
| | | Treatment | | 2 | 3.61 | | 0.035 |
| | | Residuals | | 45 | | | |
| | | | 0 vs Low | | | 0.44 | 0.659 |
| | | | 0 vs Intermediate | | | -2.07 | 0.044 |
| Colistin-Furanone C30 | | | | | | | |
| | - Atb | | 0 vs reference line | 19 | | -3.38 | 0.003 |
| | | | Low vs reference line | 15 | | -3.85 | 0.001 |
| | | | Intermediate vs reference line | 15 | | -3.20 | 0.006 |

| | | | | | | | |
|---|---|---|---|---|---|---|---|
| | + Atb | | 0 vs reference line | 19 | | 4.54 | $2.223 \times 10^{-4}$ |
| | | | Low vs reference line | 19 | | 5.30 | $4.121 \times 10^{-5}$ |
| | | | Intermediate vs reference line | 19 | | 2.48 | 0.023 |
| | | Treatment | | 2 | 13.73 | | $1.358 \times 10^{-5}$ |
| | | Residuals | | 57 | | | |
| | | | 0 vs Low | | | -4.32 | $6.320 \times 10^{-5}$ |
| | | | 0 vs Intermediate | | | -4.73 | $1.530 \times 10^{-5}$ |
| Meropenem-Furanone C30 | | | | | | | |
| | - Atb | | 0 vs reference line | 15 | | -2.62 | 0.019 |
| | | | Low vs reference line | 7 | | -1.57 | 0.159 |
| | | | Intermediate vs reference line | 7 | | -1.70 | 0.133 |
| | + Atb | | 0 vs reference line | 15 | | 5.73 | $3.949 \times 10^{-5}$ |
| | | | Low vs reference line | 15 | | 6.33 | $1.348 \times 10^{-5}$ |
| | | | Intermediate vs reference line | 15 | | 7.73 | $1.315 \times 10^{-6}$ |
| | | Treatment | | 2 | 1.15 | | 0.326 |
| | | Residuals | | 45 | | | |
| | | | 0 vs Low | | | 0.38 | 0.706 |
| | | | 0 vs Intermediate | | | 1.46 | 0.151 |
| Tobramycin-Furanone C30 | | | | | | | |
| | - Atb | | 0 vs reference line | 26 | | -12.23 | $2.732 \times 10^{-12}$ |

| | | | | | | | |
|---|---|---|---|---|---|---|---|
| | | | Low vs reference line | 15 | | -9.78 | $6.721 \times 10^{-8}$ |
| | | | Intermediate vs reference line | 15 | | -4.55 | $3.844 \times 10^{-4}$ |
| | + Atb | | 0 vs reference line | 26 | | 8.80 | $2.815 \times 10^{-9}$ |
| | | | Low vs reference line | 26 | | -3.17 | 0.004 |
| | | | Intermediate vs reference line | 26 | | -6.11 | $1.865 \times 10^{-6}$ |
| | | Treatment | | 2 | 71.61 | | $< 2.2 \times 10^{-16}$ |
| | | Residuals | | 78 | | | |
| | | | 0 vs Low | | | -9.26 | $3.390 \times 10^{-14}$ |
| | | | 0 vs Intermediate | | | -11.20 | $< 2.0 \times 10^{-16}$ |

[1]For each combination we performed a t-test. Degrees of freedom, t-values and p-values refer to the comparison to the zero line of the fitness of AtbR clones in presence of different concentrations of the antivirulence compound without the antibiotic.

[2]For each combination we performed an ANOVA. Degrees of freedom, F-values and p-values refer to the effect of the factor „Treatment“ on the fitness of AtbR clones.

**S2 Table. Concentrations of antibiotics and antivirulence drugs used for the combinatorial treatments and the competition assays**

| Compound | Media | Concentrations for combination experiments | Concentration for competitions |
|---|---|---|---|
| Ciprofloxacin | CAA+Tf | 0.003, 0.006, 0.013, 0.025, 0.031, 0.05, 0.1, 0.25 (µg/ml) | 0.013 µg/ml |
| | CAS | 0.03, 0.06, 0.13, 0.19, 0.25, 0.38, 0.5, 1 (µg/ml) | 0.25 µg/ml |
| Colistin | CAA+Tf | 0.005, 0.019, 0.023, 0.035, 0.053, 0.075, 0.15, 0.3 (µg/ml) | 0.075 µg/ml |
| | CAS | 0.16, 0.31, 0.44, 0.63, 0.66, 0.99, 1.25, 1.48 (µg/ml) | 0.99 µg/ml |
| Meropenem | CAA+Tf | 0.03, 0.06, 0.125, 0.25, 0.5, 0.875, 1.75, 4 (µg/ml) | 0.25 µg/ml |
| | CAS | 0.004, 0.008, 0.031, 0.063, 0.11, 0.22, 0.5, 1.75 (µg/ml) | 0.06 µg/ml |
| Tobramycin | CAA+Tf | 0.005, 0.019, 0.038, 0.075, 0.125, 0.15, 0.25, 0.6 (µg/ml) | 0.075 µg/ml |
| | CAS | 0.063, 0.19, 0.5, 0.75, 1, 1.5, 3, 6 (µg/ml) | 0.75 µg/ml |
| Gallium | CAA+Tf | 0.47, 0.94, 1.56, 3.13, 6.25, 12.5, 25, 50 (µM) | 1.56 µM (low), 6.25 µM (intermediate) |
| Furanone C-30 | CAS | 3.05, 6.3, 22.8, 34.2, 51.4, 97.5, 195, 390 (µM) | 6.3 µM (low), 22.8 µM (intermediate) |
